# Unified Generation of Regionalized Neural Organoids from Single-Lumen Neuroepithelium

**DOI:** 10.1101/2025.11.18.689013

**Authors:** Jyoti Rao, Zhisong He, Sebastian Loskarn, Audrey Bender, Youngmin Jo, Johanna Lückel, Martina Curcio, Irineja Cubela, J. Gray Camp, Barbara Treutlein, Matthias P. Lutolf

## Abstract

Human brain development begins with the formation of the neural tube, a neuroepithelium organized around a single continuous lumen that is patterned to specify distinct brain regions. Human pluripotent stem cell (hPSC)-derived neural organoids offer powerful models to study this process and its disruption in disease. However, most existing protocols rely on stochastic self-organization of hPSC aggregates, leading to high variability in tissue architecture and cell type composition, which limits reproducibility and fidelity to natural development. This variability has also hindered the broader adoption of brain organoid technology in translational and pharmaceutical research, where robustness and standardization are critical. Here, we present a scalable and reproducible platform that uses uniform, single-lumen neuroepithelium (SLN) as a standardized starting point for generating diverse, regionally specified neural organoids. SLNs form with high efficiency and reproducibility, can be patterned along dorso-ventral and anterior-posterior axes, and mature in suspension culture into organoids representing forebrain, midbrain, hindbrain, and neural retina. This unified and scalable approach provides a reproducible foundation for modeling human brain development, offering broad translational potential for mechanistic studies, disease modeling, and drug discovery.

## Introduction

The study of human brain development and its associated disorders is hampered by the limited accessibility of native tissue and capability for experimental manipulations. Human pluripotent stem cell (hPSC)-derived in vitro models - including neural organoids^1,2,3,4,5^, assembloids^6,7,8^ and engineered neural tube tissues^9,10^ - offer powerful opportunities to investigate the complex processes of neurodevelopment and the pathophysiology of neurological diseases.

Unguided protocols for neural organoid generation typically rely on floating hPSC aggregates that spontaneously self-organize into multiple neuroepithelial units surrounding distinct lumina^5^. These methods are highly effective at recapitulating complex, *in vivo*-like tissue architecture. Importantly, the arrangement and cell composition of neural progenitor cells within these units are critical for proper brain development^11^ and can recapitulate key features of human-specific development^12,13,14,15^. Guided organoid protocols utilize controlled signaling cues to direct regional cell fate and achieve improved reproducibility^16,17^. Other approaches focus on geometrically confining cells to mimic the architecture of the early neuroepithelium during early human brain development^9,10,18,19,20^.

Existing approaches to establish organized neuroepithelium for long-term differentiation rely on manual isolation of neuroepithelial units^21,22,23^, manual slicing of 2D layers of cells^24^ and specific platforms^10^. These approaches highlight a critical gap in the field: the lack of scalable, standardized methods for generating organized neuroepithelial precursors as a robust and unified starting point for brain region-specific organoid development.

Here, we present a scalable strategy to generate single-lumen neuroepithelium (SLNs) that supports differentiation into neural organoids guided towards multiple distinct brain regions. Building on our previously developed and now commercially available hydrogel microwell array technology^25^ we aggregate hPSCs in a high-throughput and reproducible manner, enabling spontaneous self-organization into apico-basally polarized SLNs. These tissues intrinsically adopt an anterior neural fate and undergo characteristic morphogenetic and fate changes in response to specific morphogen signals. Importantly, initiating neural organoid generation directly from SLNs improves scalability and structural fidelity, enabling robust recapitulation of the full anterior-posterior axis of the developing brain, including forebrain, midbrain, hindbrain, and retinal tissues. The extracellular matrix-dependent, tunable apical-basal polarity of the SLNs provides a modular platform for the differential organization of cells, facilitating the experimental manipulation and optimization of neural organoid morphology. Moreover, all regionally specified organoids are generated using uniform culture conditions, streamlining workflows and reducing technical barriers. We thus provide an alternative entry point into neural organoid generation that improves scalability, standardization, and structural fidelity, thereby enabling more accurate and translationally relevant models of human brain development.

## Results

### Generation of single-lumen neuroepithelium from human pluripotent stem cells

To generate SLNs in a robust, reproducible, and high-throughput manner, we employed commercially available PEG hydrogel-based microcavity plates^25^ (Figure 1a). By systematically optimizing the hPSC seeding density, microwell size, medium composition, and the timing and concentration of Matrigel addition, we established conditions that reliably produced SLNs.

**Figure 1.**
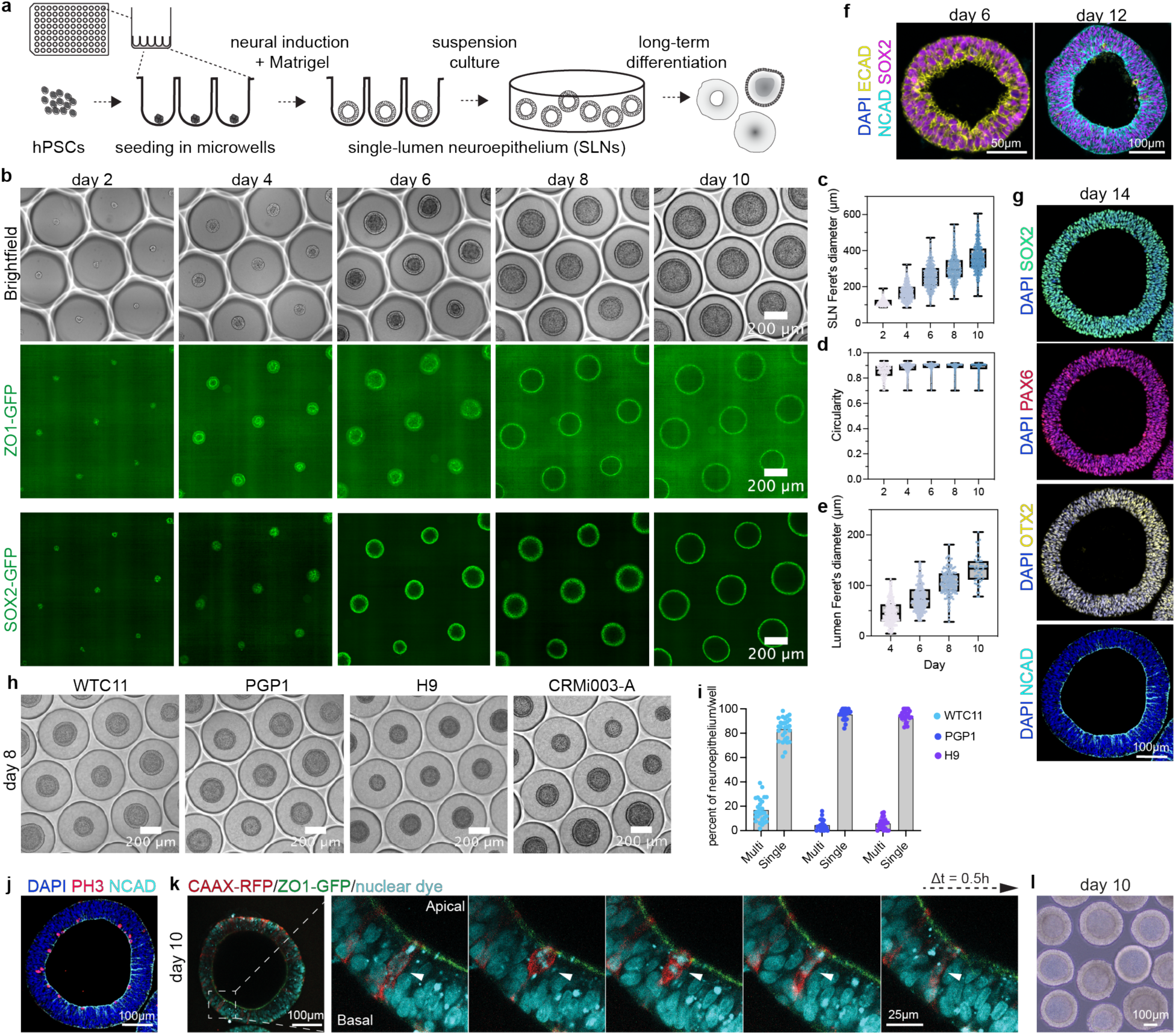
Generation of single-lumen neuroepithelium from human pluripotent stem cells. **(a)** Schematic showing the generation of SLNs in microwells and their further differentiation. (**b)** Time course imaging of developing SLNs showing the progression of differentiation and lumen formation in SLNs. **(c)** Quantification of SLN size measured using SOX2-GFP signal over time. n = 3 independent experiments. **(d)** Quantification of lumen size measured using ZO1-GFP signal over time. n = 3 independent experiments. **(e)** Quantification of SLN roundness measured using SOX2-GFP signal over time. n = 3 independent experiments. **(f)** Immunofluorescence antibody staining for E-cadherin (ECAD), N-cadherin (NCAD), and SOX2 on day 6 and day 12 of differentiation. (**g)** Immunofluorescence staining for neuroepithelial markers on day 14 of differentiation. (**h)** Representative images on day 6 of differentiation of four independent pluripotent stem cells lines including WTC-11 iPSCs (GM25256), PGP1 iPSCs (GM23338), H9 ESCs and CRMi003-A iPSCs. **(i)** Quantification of neuroepithelium consisting of single-(single) or multiple-lumina (multi) in various hPSC lines. n = 3 independent experiments (**j)** Immunofluorescence antibody staining for phospho-histone 3 (PH3) and apical marker NCAD in the SLNs. (**k)** Live imaging snapshots of a SLN derived from a mix of ZO1-GFP WTC-11 and CAAX-RFP WTC-11 iPSCs. Arrow indicates the dividing cells at the apical surface and exhibiting interkinetic nuclear migration. **(l)** 10-day-old SLNs transferred out of microwell into an ultra-low attachment plate for further differentiation.

Reducing the seeding cell number for unguided organoid differentiation^5,26^ resulted in fewer neuroepithelia in the organoids. Very rare SLNs could be observed in the 10-cells condition during early days of differentiation which matured into multi-neuroepithelial structures over time (Supplementary Figure 1a). To provide physical confinement for better self-organization and higher efficiency SLN formation, we tested various microwell sizes. A 500 μm diameter microwell was optimal for SLN generation, whereas larger microwells or standard 96-well plates did not support single-neuroepithelium formation or stability, while smaller wells restricted growth (Supplementary Figure 1b-c). Additionally, Matrigel concentration was critical: low concentrations (2.5%) resulted in thin neuroepithelium, whereas 5% to 20% Matrigel generated uniform SLN morphogenesis (Supplementary Figure 1d).

SLN differentiations were monitored using GFP-tagged SOX2 and ZO1 reporter iPSC lines. SLN consisted of SOX2-positive epiblast-like cells with apical ZO1 expression (Figure 1b), grew steadily in size and lumen diameter while maintaining a high circularity (Figure 1c-e). Immunofluorescence confirmed a progressive developmental transition: expression of the epiblast marker E-cadherin at day 6, replacement by the neural marker N-cadherin at day 12 (Figure 1f), and maintenance of structural integrity alongside neural marker expression (SOX2, PAX6, OTX2) at day 14 (Figure 1g).

The platform was highly reproducible: four distinct hPSC lines formed SLNs with comparable morphology and efficiency (83-95%), dependent on the specific hPSC cell line used (Figure 1h-i). To dissect the reason for multi-lumen formation, we tracked initial cell distribution and final SLN phenotype (Supplementary Figure 1e). Uneven cell distribution within the microwell, likely due to surface tension, resulted in a higher density of cells near the well edges (Supplementary Figure 1f). Corroborating this, higher initial cell numbers correlated with multi-lumen formation (Supplementary Figure 1g-1h). Of note, SLNs could also be generated in alternative microwell formats (Elplasia and Aggrewell plates), indicating that confinement, rather than substrate material, is the key determinant of organized aggregation, although subtle size differences were observed between formats (Supplementary Figure 1i–k).

Dividing cells in late G2/M phase (phospho-histone 3 positive) were consistently observed at the apical surface, suggesting interkinetic nuclear migration (Figure 1j). Live imaging of mosaic SLNs (sparsely labelled CAAX-RFP iPSCs hashed into the ZO1-GFP iPSCs) revealed nuclei migrating apically to divide at the ventricular-like surface of the pseudostratified epithelium (Figure 1k, Supplementary Figure 1l). After day 10, SLNs were easily transferred to ultra-low attachment plates for long-term culture (Figure 1l). Finally, applying this protocol to mouse embryonic stem cells also yielded SLNs (Supplementary Figure 1m), indicating that the principles of in vitro SLN formation are species-independent and broadly applicable.

### SLNs adopt a default anterior state but retain regional plasticity

To characterize the cellular composition of SLNs, we performed single-cell RNA-sequencing (scRNA-seq) at day 10. SLNs displayed a high degree of transcriptional homogeneity, with variation mainly reflecting cell cycle states, ribosomal protein expression, and detection rates (Figure 2a, Supplementary Figure 2a). Cells expressed canonical neuroepithelial markers (NES, MKI67) together with the anterior neural marker OTX2 but lacked HOX genes (HOXA2, HOXB13) and posterior marker WLS (Figure 2b), indicating a default anterior neural identity. When dissociated and reaggregated, day 10 SLNs rapidly re-formed lumenized structures (Supplementary Figure 2b), confirming their identity as polarized neuroepithelial progenitors.

**Figure 2.**
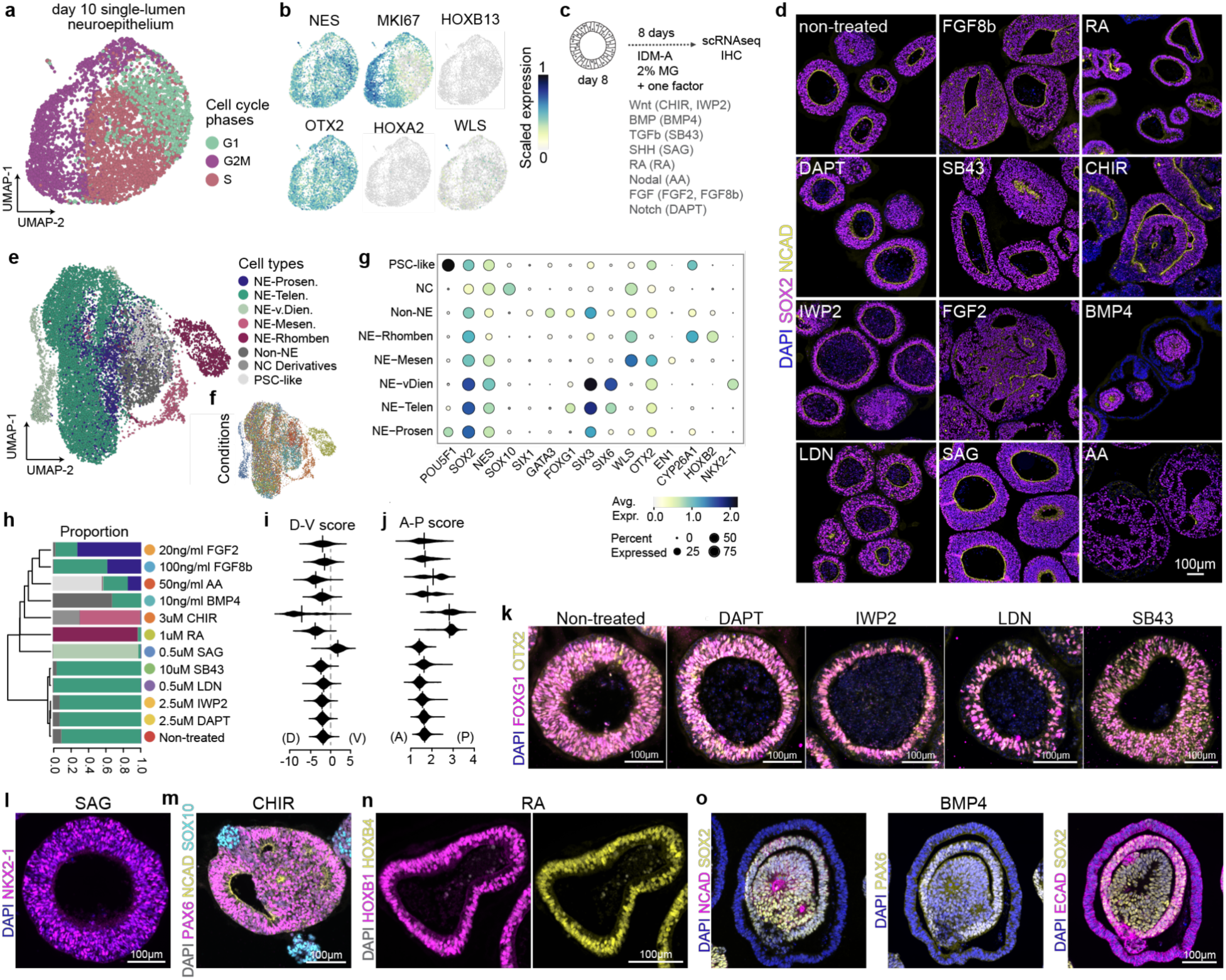
SLNs adopt a default anterior state but retain regional plasticity. **(a)** UMAP visualization of single cell transcriptomics data from day 10 SLNs, colored according to their inferred cell cycle phases (G1, S, G2M). **(b)** Feature plots overlaid on the UMAP shown in (a). **(c)** Schematic overview of the morphogen treatment and downstream analysis. Multiplexed single cell analysis was performed on multiple pooled organoids. scRNAseq - single cell RNA sequencing, IHC - immunohistochemistry. **(d)** Representative immunofluorescence images showing the morphological changes and expression of SOX2 and N cadherin. Nuclei are counterstained with DAPI. SB43 - SB431542, LDN - LDN193189CHIR - CHIR99021, RA - retinoic acid, AA - Activin A. **(e)** UMAP embedding of the integrated single-cell transcriptomes from all treatment conditions, with cells colored by cell type annotation derived from Braun et al.^27^. NE- neuroepithelium, Prosen - Prosencephalic, Telen - Telencephalic, vDien- ventral Diencephalic, Mesen - Mesencephalic, Rhomben - Rhombencephalic, Non-NE - non-neuroepithelium, NC - neural crest, PSC-like - pluripotent stem cel-likel. **(f)** UMAP embedding as in (e) with cells colored according to the different morphogen treatment groups. **(g)** Dot plot showing the expression levels and percentage of expressing cells for selected regional identity marker genes across the different cell type annotations. **(h)** Hierarchical clustering of transcriptomics data and bar graph showing the proportion of different cell types under various differentiation conditions. **(i)** Violin plots illustrating the dorso-ventral (D-V) patterning score of SLNs under treatment conditions. **(j)** Violin plots illustrating the anterior-posterior (A-P) patterning score of SLNs under treatment conditions. Immunofluorescence analysis of SLNs **(k)** for DAPI (nucleus) and FOXG1 (dorsal forebrain) after anteriorizing morphogen treatment **(l)** for ventral marker NKX2-1 after SAG treatment **(m)** for neuroepithelial marker PAX6 and neural crest cells marked by SOX10 after CHIR treatment **(n)** for HOX genes after RA treatment **(o)** for PAX6, SOX2, neuroepithelial apical marker NCAD and non-neural epithelial marker ECAD after BMP4 treatment.

We next examined how developmental signaling pathways influenced SLN fate and morphology (Figure 2c). SLNs were treated with modulators of different signaling pathways on day 8 of differentiation. After 8 days of treatment, the SLNs were harvested for immunostaining and scRNAseq analysis (Supplementary Figure 2c). The SLNs showed pronounced morphological responses in response to the treatment (Figure 2d). Untreated SLNs, or those exposed to anteriorizing inhibitors of Wnt (IWP2), TGFβ (SB431542), BMP (LDN193189), or Notch (DAPT), retained a simple spherical morphology (Figure 2d, Supplementary Figure 2c). In contrast, activation of Wnt (CHIR99021), BMP (BMP4), retinoic acid (RA), FGF (FGF2, FGF8), or Nodal (Activin A) pathways induced a drastic alteration in tissue organization (Figure 2d, Supplementary Figure 2c).

To assess regional identity, we performed multiplexed scRNA-seq of treated organoids (approximately 100 pooled organoids per condition) (Figure 2c). We projected the processed data onto a reference atlas of first-trimester developing human brains^27^ using reference similarity spectrum analysis^28^. UMAP embedding revealed cell populations corresponding to prosencephalic (telencephalic and diencephalic), mesencephalic, and rhombencephalic populations, consistent with regionalization of the developing brain. Additionally, non-neuroepithelial cell populations, including non-neural ectoderm, neural crest derivatives and PSC-like populations, were identified (Figure 2e-g).

Quantification of dorso-ventral (DV) and anterior-posterior (AP) scores showed that Wnt, TGFβ, BMP, and Notch inhibition - or no treatment - consistently yielded telencephalic progenitors (Figure 2h) robustly expressing FOXG1 (Supplementary Figure 2d–e), indicating maintenance of a dorsal-anterior identity (Figure 2i–j). All anteriorizing conditions preserved spherical morphology, suggesting that suppression of posteriorizing influences stabilizes a uniform state (Figure 2k, Supplementary Figure 2f). Shh activation (SAG) produced progenitors with a ventral-anterior identity expressing NKX2-1 (Figure 2l, Supplementary Figure 2d). Consistent with their anterior identity, SAG-treated SLNs also maintained SLN morphology (Figure 2l, Supplementary Figure 2c).

Activation of Wnt and retinoic acid signaling promoted posterior tissue fates and led to a drastic morphology change (Figure 2d, 2h, 2j, Supplementary figure 2c). Wnt activation produced mesencephalic progenitor cells (OTX2- and EN2-positive) and SOX10-positive neural crest derivatives (Figure 2m, Supplementary Figure 2d, 2g). RA activation drove the cells into rhombencephalic fate with expression of CYP26A1 and HOX genes including HOXB1 and HOXB4 (Figure 2n, Supplementary Figure 2d). BMP induced multilayered structures resembling early neural tube morphogenesis^20^, with a mix of non-neural ectoderm and telencephalic progenitors (Figure 2o, Supplementary Figure 2i-k). The inner layer formed a PAX6- and FOXG1-positive anterior neuroepithelium lined apically by NCAD (Figure 2o, Supplementary Figure 2l). The intermediate non-neural layer expressed SOX2, FOXG1 and E cadherin suggesting formation of surface ectoderm (Figure 2o, Supplementary Figure 2h). Finally, E cadherin-positive, FOXG1-negative cells expressing ectoderm markers such as GATA3 and ISL1 formed the outermost cell layer (Figure 2o, Supplementary Figure 2h, 2l).

FGF (FGF2, FGF8b) treatment dissolved SLN architecture, drove rapid proliferation, and generated mixed populations of FOXG1-positive telencephalic progenitors, POU5F1-positive PSCs, and posterior CYP26A1-positive cells (Supplementary Figure 2d–e, 2m). Activin A maintained POU5F1 expression and largely preserved PSC identity (Supplementary Figure 2d–e, 2n), consistent with its limited role in early neural tube patterning.

Collectively, these results show that SLNs adopt a default anterior fate but retain plasticity, acquiring diverse regional identities and morphologies along DV and AP axes in response to morphogen cues. This tractable system thus provides a powerful platform to dissect how signaling pathways orchestrate early human brain regionalization and tissue morphogenesis.

### Generation of regionally specified brain tissues from SLNs

We next tested whether SLNs could be cultured long-term into distinct regionalized brain organoids. Building on established protocols for forebrain^4,17,29^, midbrain^29,30,31^ and hindbrain organoids^32,33,31^, we optimized the timing and duration of morphogen treatments applied to SLNs (Figure 3a).

**Figure 3.**
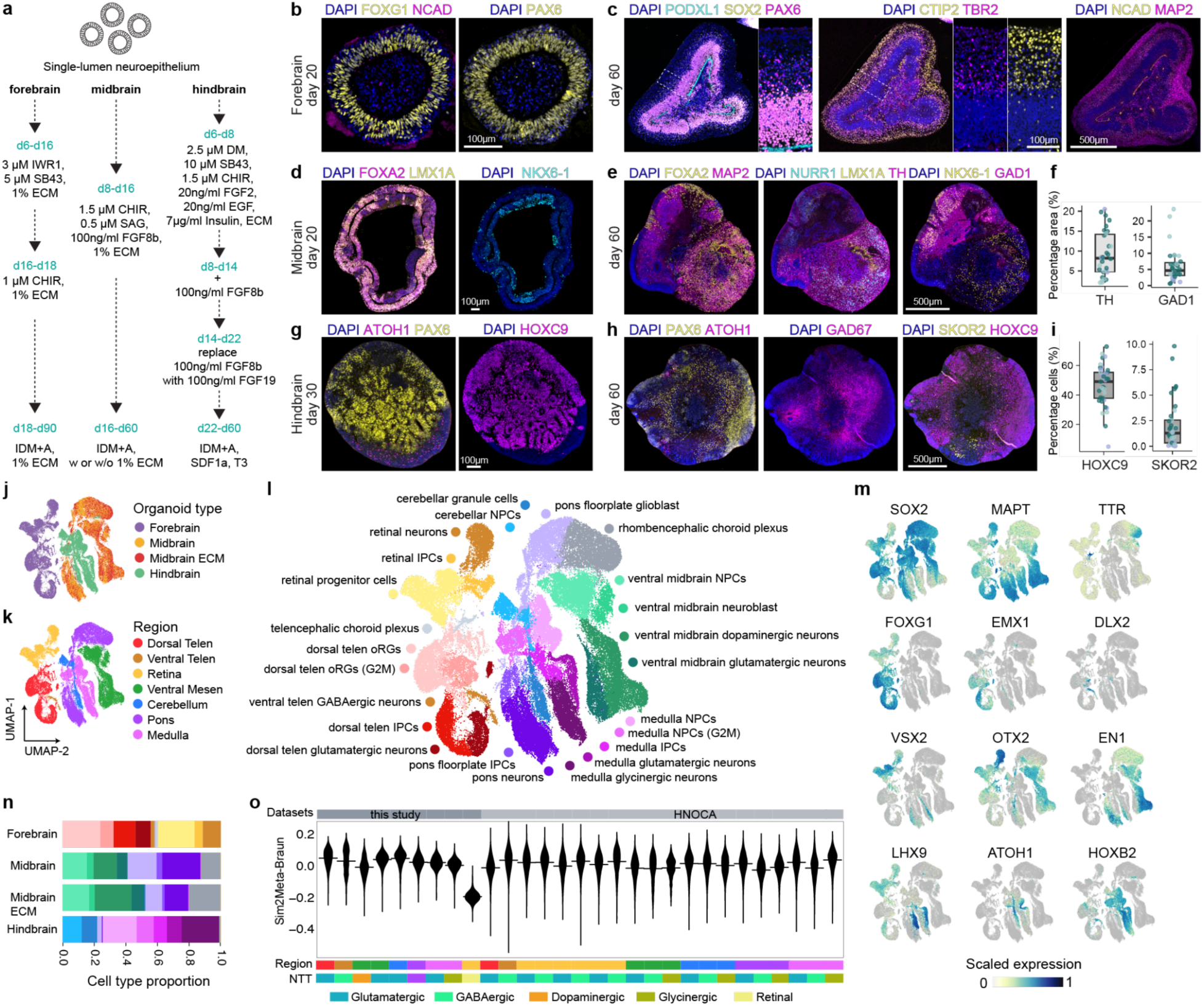
Generation of regionally specified brain organoids from SLNs. **(a)** Schematic diagram illustrating the differentiation strategy to generate forebrain, midbrain, and hindbrain organoids from SLNs using specific combinations of growth factors. **(b)** Immunohistochemistry analysis of a day 20 forebrain organoid section derived from SLN, demonstrating the expression of the dorsal forebrain progenitor marker FOXG1 and the apical polarity marker NCAD. Nuclei are counterstained with DAPI. **(c)** Immunohistochemistry analysis of a day 60 forebrain organoid, revealing distinct populations of forebrain progenitors (SOX2, FOXG1), intermediate progenitors (TBR2), and neurons (CTIP2, MAP2). Magnified views show organisation of progenitors (SOX2, TBR2) and neurons (CTIP2) in the developing organoids. **(d)** Immunofluorescence images of a day 20 midbrain organoid section stained with DAPI and midbrain progenitors marked by FOXA2, LMX1a and NKX6-1. **(e)** Immunofluorescence images of a day 60 midbrain organoid section stained with DAPI, TH (dopaminergic neurons), FOXA2, LMX1A, NURR1 (dopaminergic neuron precursors) and GABAergic neuron marker GAD67. **(f)** Quantification of TH and GAD67 positive neurons on day 60 (30 organoids from 3 independent replicates). **(g)** Immunofluorescence images of a day 30 hindbrain organoid section stained with DAPI, ATOH1, PAX6 and HOXC9. **(h)** Immunofluorescence images of a day 60 hindbrain organoid section stained with progenitors (PAX6, ATOH1), GABAergic neurons (GAD67), SKOR2 (Purkinje cell) and posterior marker (HOXC9). **(i)** Quantification of HOXC9 and SKOR2 on day 60 (37 organoids from 3 independent replicates). **(j-l)** UMAP of regional organoid transcriptomes based on gene expression profiles, colored by annotation of organoid protocol type, region and identified cell type. **(m)** Feature plots of gene expression levels of selected cell type markers across different cell types. **(n)** Proportions of cells assigned to different cell types based on reference projection. **(o)** Spearman correlations between gene expression profiles of neural cell types in organoids and HNOCA datasets and those in the human developing brain atlas.

#### Forebrain organoids

Soluble extracellular matrix (ECM, 1% Matrigel) was essential to preserve apical–basal polarity and its removal caused inversion of polarity and a disrupted architecture (Supplementary Figure 3a), underscoring the role of ECM in maintaining neuroepithelial organization^34^. SLNs treated with 5μM TGFβ inhibitor SB431542 and 3μM Wnt inhibitor IWR1^4,17^ from day 6-16, followed by Wnt activation with 1μM CHIR99021 (day 16-18) in presence of soluble ECM, efficiently adopted forebrain identity (Figure 3a, Supplementary Figure 3b). By day 30, organoids expressed the anterior–dorsal marker FOXG1 and mostly contained a single N-cadherin lined neuroepithelial unit (Figure 3b, Supplementary Figure 3c). By day 60, the structural complexity increased and multiple neuroepithelial units emerged in the organoids (Figure 3c, Supplementary Figure 3d). The organoids expressed canonical cortical markers PAX6, TBR2 and CTIP2 (Figure 3c). In addition, several organoids consisted of retinal neuroepithelium marked by VSX2 (Supplementary Figure 3d).

#### Midbrain organoids

Treatment with 1.5 μM CHIR99021, the Shh agonist 0.5 μM SAG, and 100 ng/ml FGF8b^29,30,35,31^ from day 8-16 in presence of soluble ECM (1% Matrigel) induced rapid epithelial expansion and complex morphologies (Figure 3a, Supplementary Figure 3e-f). By day 30, organoids consisted of polarized extended neuroepithelium expressing ventral midbrain progenitor markers FOXA2, LMX1A, and NKX6-1 (Figure 3d, Supplementary Figure 3f-g). Over time, the organoids continued to maintain ventral midbrain progenitors in addition to dopaminergic neurons (TH) and GABAergic neurons (GAD67) (Figure 3e–f, 3h–i). Presence of soluble ECM at later stages increased tissue size and structural complexity but not overall fate specification, although it resulted in increased variability in cell type composition (Supplementary Figure 3f, 3h-i).

#### Hindbrain organoids

SLNs were exposed to dual SMAD inhibition (2.5 μM Dorsomorphin, 10 μM SB431542) together with posteriorizing factors (1.5 μM CHIR99021, 20ng/mL FGF2, 20ng/mL EGF, 7 μg/ml insulin)^32,33^ from day 6-8 (Figure 3a, Supplementary Figure 4a). From day 8-14, 100 ng/ml FGF8b was added to the first medium. From day 14-22 FGF8b was replaced by FGF19. By day 30, organoids formed multiple neuroepithelial units consisting of SOX2-positive progenitors (Supplementary Figure 4b). Distinct populations emerged, including mutually exclusive PAX6-positive cells and ATOH1-positive rhombic lip progenitors. Notably, ATOH1-positive cells lacked HOXC7, suggesting representation of multiple hindbrain rhombomeres (Figure 3g, Supplementary Figure 4c–d). By day 60, SKOR2-positive Purkinje cells localized adjacent to ATOH1-positive germinal cells, alongside GABAergic neurons and HOXC7-positive posterior cells (Figure 3h–i, Supplementary Figure 4e–f).

To profile the cell types generated in the regionally specified organoids, we generated single cell transcriptomics data from 90-day old forebrain and 60-day old midbrain and hindbrain organoids. Integrated scRNA-seq data revealed that cells from each regional organoid type clustered distinctly, corresponding directly to the *a priori* defined protocols (Figure 3j, 3k). Forebrain organoids were primarily composed of cells mapping to the dorsal (FOXG1 and EMX1) and ventral (DLX2) telencephalon regions, with cell types including dorsal telencephalic neural progenitors, intermediate progenitor cells, dorsal glutamatergic (SLC17A7) and ventral GABAergic neurons (DLX2, GAD2), as well as some telencephalic choroid plexus and retinal progenitors (VSX2) (Figure 3l-n, Supplementary Figure 5a). Midbrain organoids were dominated by cell types characteristic of the ventral mesencephalon (OTX2 and EN1), including ventral midbrain neural progenitors and neuroblasts (LMX1A, FOXA2, EN1), and both dopaminergic (TH) and glutamatergic (SLC17A6) neurons. Long-term ECM exposure did not have a major impact on the cellular composition of the midbrain organoids (Figure 3n). Notably, midbrain organoids also contained a significant portion of pons cell types including pons floorplate progenitors (FOXA2) and neurons (SLC17A6), indicating some caudal regional spread beyond the midbrain designation (Figure 3l-n, Supplementary Figure 5a). Hindbrain organoids showed a strong composition of cells mapping to the cerebellum and medulla regions (LHX9, ATOH1, and HOXB2). Specific cell types include cerebellar progenitors and granule cell neurons (LHX9), medulla neural progenitors (HOXB2) and medulla glycinergic neurons (SLC6A5) (Figure 3l-n, Supplementary Figure 5a).

To validate the biological relevance of SLN-derived organoids, we compared the regional organoid data to established human brain reference atlases. Hierarchical clustering of our single-cell data with the Human Neural Organoid Cell Atlas (HNOCA) dataset^28^ demonstrated that SLN-derived organoids robustly clustered with their corresponding HNOCA counterparts (Supplementary Figure 5b). Specifically, hindbrain organoids clustered with a dataset focused on human spinal cord and hindbrain development^36^, while midbrain organoids clustered with a dataset focused on ventral midbrain organoids^35^. Furthermore, we assessed the transcriptomic fidelity of the neuronal cell types by comparing them to the primary reference atlas from Braun et al.^27^. SLN-derived regional organoids generated neuronal cell types with comparable similarity to the corresponding primary reference cell types as HNOCA datasets, highlighting their accurate molecular identity (Figure 3o). These analyses provide evidence that SLN-derived organoids faithfully recapitulate the transcriptional trajectories of conventional organoids.

### Scalable retinal organoids with ECM-tunable polarity

Current protocols for generating retinal organoids from hPSCs often rely on excising neural retina from mixed 3D tissues^37^ or on intermediate 2D culture steps^38,39,40,41^. Fully suspension-based methods have also been described^42,43^, but a persistent challenge across approaches is the mislocalized co-generation of retinal pigmented epithelium (RPE) alongside neural retina. To address this, we adapted the SLN platform to a retinal-specific differentiation paradigm analogous to that used for brain regionalization (Figure 3a).

SLNs were directed into early eye fields by inhibiting TGFβ (SB431542), BMP (Dorsomorphin), and Wnt (IWR1) signaling^37,44^, followed by BMP4 treatment to expand retinal progenitors^42^ and long-term maturation media^40^ (Figure 4a). We leveraged the structural malleability of SLNs to determine the effect of ECM on retinal organoid polarity. Presence of soluble ECM (1% Cultrex) in the medium dictated both the final size and the apical-basal orientation of the developing retinal epithelium (Figure 4b-c). Retinal fate commitment was confirmed across both conditions by the expression of retinal progenitor marker VSX2 and photoreceptor progenitor marker CRX. Organoids differentiated without ECM displayed an apical-out configuration, with a majority of CRX-positive cells localizing to the exterior surface. Conversely, ECM-treated organoids adopted an apical-in polarity, positioning the CRX-positive cells layer internally towards the lumen (Figure 4d). This demonstrates that ECM modulation provides a mechanism for manipulating the structural orientation of retinal tissues.

**Figure 4.**
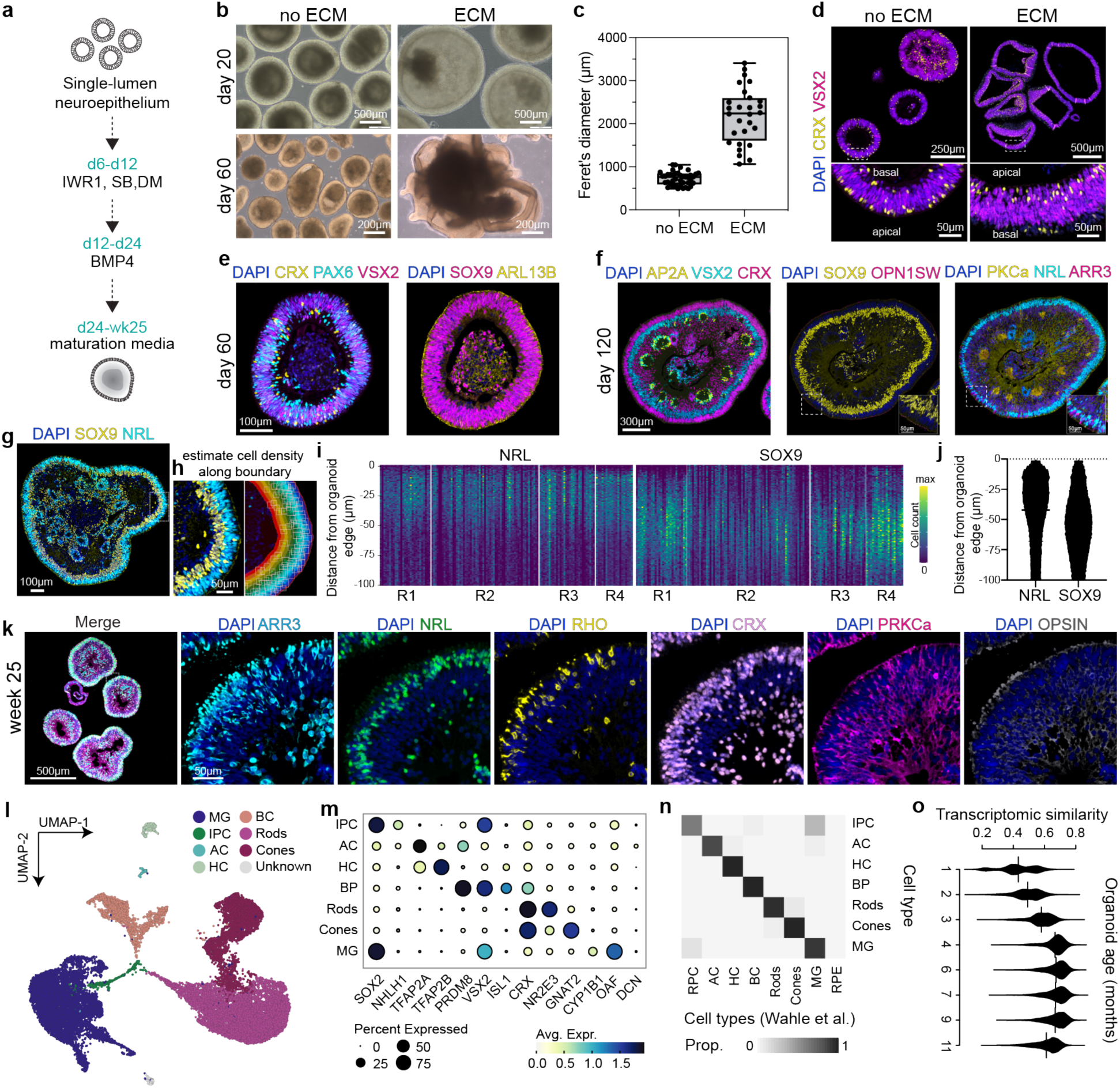
Scalable retinal organoids with ECM-tunable polarity. **(a)** Schematic illustrating the differentiation of SLNs into retinal organoids in fully suspension culture. **(b)** Brightfield images of retinal organoids cultured with or without ECM (1% Cultrex) at day 20 and day 60. **(c)** Quantification of organoid size on day 60. (n=2 independent experiments). **(d)** Immunofluorescence images of day 60 retinal organoids cultured with or without ECM. Inset shows magnified view with apical and basal compartment. **(e)** Immunofluorescence images of a day 60 no ECM retinal organoid section stained with DAPI (nuclei) and antibodies against retinal progenitor markers PAX6, VSX2 and SOX9, and photoreceptor progenitor marker CRX. **(f)** Immunofluorescence staining of no ECM retinal organoids at day 120, showing the expression of various retinal cell type markers, including photoreceptor progenitor cells (CRX), photoreceptors (OPN1SW, NRL, ARR3), bipolar cells (PKC-a), Amacrine cells (AP2A), Müller glia (SOX9). Insets show magnified views of specific regions. **(g)** Immunofluorescence staining of retinal organoids on day 120, demonstrating the expression and localization of NRL (outer nuclear layer) and SOX9 (inner nuclear layer). **(h)** Magnified view of the highlighted region from (g), showing counted cells (boxed) and the concentric bands used for spatial analysis. **(i)** Heatmaps illustrating the spatial distribution of SOX9 and NRL positive cells as a function of distance from the organoid edge, suggesting the formation of distinct retinal layers. Color intensity indicates cell count. (n= 4 independent experiments,146 organoids measured). **(j)** Scatter plot showing overall distribution of NRL and SOX2 positive cells measured in (h) at the organoid edge. (n=4 independent experiments). **(k)** Immunofluorescence staining of retinal organoids at week 25, showing the mature expression of various retinal cell type markers, including ARR3 (cones), NRL (rods), RHO (rod opsin), PRKCa (bipolar cells), and Opsin (photoreceptors). **(l)** UMAP visualization of single cells from week 25 retinal organoids, colored by major retinal cell types (MG: Müller glia, IPC: Intermediate Progenitor Cells, AC: Amacrine Cells, HC: Horizontal Cells, BC: Bipolar Cells, Rods, Cones, ECM: Extracellular Matrix expressing). **(m)** Dot plot showing the expression levels and percentage of expressing cells for selected marker genes across the identified cell types. **(n)** Heatmap displaying proportion of cell type label matching between the cell types identified in SLN-derived organoids and previously defined cell types in published retinal organoids^45^. **(o)** Violin plots showing transcriptomic similarity scores of organoids generated in this study to organoids across different time points from a published time-course retinal organoid dataset^45^.

Organoids without ECM transitioned from cystic neuroepithelium to structured retinal tissues with photoreceptor outer segments by week 30 (Supplementary Figure 6a). Size increased until ∼day 90, after which it plateaued (Supplementary Figure 6c). Morphology-based selection at day 120 enriched for well-formed retinal organoids (Supplementary Figure 6b). By day 60, organoids expressed early retinal progenitor markers PAX6, VSX2, and SOX9, along with the photoreceptor precursor CRX (Figure 4e). By day 120, they contained multiple retinal cell types, including CRX-positive progenitors, mature photoreceptors (ARR3, OPN1SW for cones; NRL for rods), PKC-α-positive bipolar cells, AP2α-positive amacrine cells, and SOX9-positive Müller glia (Figure 4f). To quantify the organization of outer and inner nuclear layers, we analyzed NRL-positive outer nuclear layer and SOX9-positive Müller glia and other inner nuclear layer cells distribution in organoids embedded in tissue microarrays on day 120 (Figure 4g, Supplementary Figure 6d). We profiled cellular localization within 100 μm inward from the organoid’s periphery across over 150 organoids from four independent experiments (Figure 4h). This analysis confirmed the consistent formation of distinct, stratified layers in most organoids, with NRL-positive cells predominantly forming an outer layer and SOX9-positive cells enriched in a more inner region (Figure 4i-j).

At week 25, organoids exhibited advanced laminar structure and mature marker expression (Figure 4k). scRNA-seq profiling at week 25 revealed all major retinal cell types, including Müller glia, intermediate progenitors, amacrine cells, horizontal cells, bipolar cells, rods, and cones (Figure 4l–m). Marker gene analysis confirmed lineage identities - for example, rods expressed NRL and NR2E3, cones expressed GNAT2 and OPN1SW, and Müller glia expressed SOX9 and DCN (Figure 4m, Supplementary Figure 6e). To benchmark against existing methods, we compared the transcriptomics data from SLN-derived retinal organoids with published datasets^45^. Correlation analysis showed strong similarity of the SLN-derived retinal cell types with those in established protocols (Figure 4n). Maturation trajectory mapping further indicated that SLN-derived week-25 organoids most closely resembled ∼6-month-old reference organoids (Figure 4o).

In summary, we successfully adapted the SLN platform to develop an efficient and scalable method for generating retinal organoids. This approach overcomes existing challenges in reproducibility and polarity control, enabling the generation of highly organized retinal tissues entirely in suspension culture. By tuning the ECM environment, apical–basal polarity can be precisely modulated to direct tissue orientation, providing experimental access to both apical-in and apical-out configurations. These features not only streamline retinal organoid production but also open new opportunities to model human retinal development, maturation, and disease progression with unprecedented structural precision. More generally, the ability to control tissue polarity and organization within a unified SLN-based framework extends the versatility of this system beyond brain regionalization, establishing a scalable foundation for comparative studies of neuroepithelial morphogenesis and translational applications in vision research.

## Discussion

Here we present a user-friendly and reproducible platform for generating uniform SLNs from hPSCs and demonstrate their versatility as a precursor for diverse regionally specified brain organoids. A central feature of this approach is the precise physical confinement of a small number of cells in hydrogel^25^ or ULA-bottom microwell plates, combined with soluble Matrigel to promote polarization and self-organization. This controlled environment reproducibly directs hPSCs to form single-lumen epiblast cysts that differentiate into polarized neuroepithelial structures (Figure 1a). Compared with strategies that require manual excision of individual units^21,22,24,23^ or rely on highly specialized bioengineering approaches^10^, the presented method is scalable, accessible, and less labor-intensive. Importantly, the use of soluble rather than solid (*i.e.*, undiluted) ECM facilitates long-term culture while minimizing manual handling and increasing scalability. By systematically optimizing microwell geometry, ECM composition, and input cell number, we significantly improved SLN formation efficiency and reproducibility. Robust derivation of SLNs across multiple hPSC lines (WTC-11, PGP1, H9, CRMi003-A) and mouse ESCs further highlights the generalizability of this principle across hPSC sources and even species.

scRNA-seq analysis confirmed that SLNs represent a pure neuroepithelial population, unlike hPSC 3D aggregates based methods, which generate a mixed composition including non-neural lineages^46^. SLNs maintained key hallmarks of early neuroepithelium - apicobasal polarity and interkinetic nuclear migration - and exhibited an intrinsic anterior bias (OTX2-positive, HOX genes and WLS-negative). Thus, SLNs provide a highly defined and consistent starting point for directed differentiation, enabling more controlled studies of patterning and morphodynamics in early brain development.

Recent multiplexed morphogen screens have mapped signaling logic for neural regionalization^31,47,48^. Our work complements and extends this framework by showing that SLNs are highly responsive to developmental cues, including Wnt, BMP, retinoic acid, Notch, Shh, FGF, TGFβ, and Activin. Guided by these pathways, SLNs could be robustly directed to distinct regional identities along both anterior-posterior and dorso-ventral axes. In addition, the small size, robust architectural uniformity, and single-neuroepithelium structure of the SLNs provide opportunities for precisely studying the morphogenetic changes and self-organization dynamics of neural tissue in response to specific growth factors and environmental cues.

Using SLNs as a starting point, we established reproducible protocols for generating forebrain, midbrain, hindbrain, and retinal organoids by maintaining consistent culture conditions and basal medium across different regions. We found that continuous exposure to soluble ECM was critical for forebrain organoid polarity, consistent with prior studies of ECM requirements^34,49^. Interestingly, while forebrain organoids initially preserved a single neuroepithelial unit, they transitioned to multi-unit architectures over time, suggesting that additional features such as ventricular pressure (normally provided by cerebrospinal fluid in vivo^50^) may be necessary to maintain a ventricle-like cavity. By contrast, midbrain and hindbrain organoids underwent early complex morphogenetic remodeling.

scRNAseq analysis confirmed that forebrain organoids faithfully formed dorsal and ventral telencephalon cell types, including retinal cells. Midbrain and hindbrain organoids generated their target lineages (ventral mesencephalon and cerebellum/medulla, respectively), but exhibited regional overlap, most notably the presence of pons cells in the midbrain cultures. This suggests a further refinement of patterning signaling factors is needed to generate purer midbrain organoids^51^. Single-cell transcriptomics also confirmed high similarity of SLN-derived organoid cell populations to those in published reference atlases^28^, validating their developmental fidelity.

A notable advance of the presented system is a simplified method for the generation of retinal organoids entirely in suspension without ectopic RPE formation^52,39,40,41^. SLN-derived retinal organoids exhibited progressive morphogenesis, robust laminar organization, and a full repertoire of retinal cell types^45^. scRNA-seq at week 25 confirmed advanced maturation, with transcriptomes strongly correlating with those of published late-stage organoids^45^. Interestingly, SLN-derived retinal tissues exhibit an ECM-tunable apical-basal polarity. This control is crucial as it could potentially facilitate the organized maturation and accessibility of retinal ganglion cells (RGCs) on the organoid surface, overcoming the survival challenge often faced when these cells reside on the inaccessible inner side^45,52^. The preservation and accessibility of RGCs are particularly important, as these neurons form the output layer of the retina and transmit visual information to the brain. Their maintenance in vitro would not only enhance the functional completeness of retinal organoids but also broaden their utility for modeling optic neuropathies and screening neuroprotective compounds.

In summary, our work establishes a highly reproducible and scalable platform for generating SLNs that serve as a standardized and uniform precursor for diverse regionalized neural organoids. This approach reliably yields forebrain, midbrain, hindbrain, and retinal tissues with high cellular and transcriptional fidelity, validating its robust capacity to recapitulate major aspects of early human brain development. By providing a simplified and unified entry point for long-term organoid culture, this platform enhances the utility of neural organoids for mechanistic studies of morphogenesis and expands their translational potential for disease modeling, regenerative medicine, and drug discovery.

## Methods

### Pluripotent stem cell culture

WTC11 (GM25256, wildtype) human induced pluripotent stem cell (iPSC) line, PGP1 (GM23338, wildtype) human iPSC line, CRMi (CRMi003-A, wildtype) and H9 (WA09, wildtype) human embryonic stem cell (ESC) line were routinely cultured on Geltrex-coated culture plates (Thermo Fisher, A1413201) in mTeSR Plus medium (Stemcell Technologies, 100-0276) at 37°C under normoxic conditions. Upon reaching confluency, cultures were dissociated into single cells using Accutase (Corning, 25-058-CI). Cells were then seeded onto Geltrex-coated plates in mTeSR Plus supplemented with 10 μM Y-27632 dihydrochloride (Tocris, 1254) at a density of 20,000-25,000 cells/cm² for routine maintenance of iPSC and ESC lines. The following day, the medium was replaced with mTeSR Plus without supplements, and medium changes occurred daily. Cultures typically reached confluency every four days and were passaged using the same dissociation and seeding protocol. For cryopreservation, cultures were dissociated into single cells using Accutase and frozen in Stem Cell Banker Medium (AMSbio, 11924). Cells were tested for mycoplasma contamination before first stocking. WTC11 (GM25256, wildtype), CAAX-RFP, ZO1-GFP, SOX2-GFP reporter cell lines and PGP1 (GM23338) were obtained from the NIGMS Human Genetic Cell Repository at the Coriell Institute for Medical Research. CRMi003-A iPSCs (NINDS material ID - ND50028) were obtained from NHCDR.

### Differentiation of pluripotent stem cells into SLNs

iPSCs/ESC cultures were dissociated into single cell suspension using Accutase. For generating SLNs, cells were plated into different microwell plate types including PEG based microwell plates (Sunbiosciences or Insphero, Gri3D® hydrogel microcavity plates), Elplasia (Corning, 4442) and Aggrewell 400 (Stem cell technologies, 34415, 34811). For a typical experiment, cells were seeded in a 400 - 500 um microwell Grid3D plate, at a density of 10-12 cells per microwell in mTeSR plus supplemented with 1:1000 CEPT cocktail^53^ [Polyamine (Sigma, P8483), Chroman1 (Medchem Express, HY-15392), Emricasan (Selleckchem, S7775), trans-ISRIB (Thermo Fisher scientific, 5284)]. On day 2, medium was replaced with a NIM medium^26^ [N2 medium - DMEM/F12 HEPES glutamine (Thermo Fisher Scientific, 11330057 + Penicillin/Streptomycin (Thermo Fisher, P0781) + 1x N2 supplement (Thermo Fisher, 17502048) + 1x MEM NEAA (Thermo Fisher, 11140035), 1ug/ml Heparin (Sigma-Aldrich, H3149-25KU]. On day 4, medium was replaced by N2 medium supplemented with 5% Matrigel. SLNs were kept in the microwell plate until day 8 or 10 in N2 medium supplemented with 5% Matrigel, with medium changes every other day.

For long term culture, SLNs were pipetted out of the microwells using a 1ml pipette and then moved to ultra low attachment 6 cm or 9cm dish (Corning, LSBio) in IDM-A medium^26^ [IDM-A medium - 50% DMEM/F12 HEPES glutamine (Thermo Fisher Scientific, 11330057 + 50% Neurobasal (Thermo Fisher, 21103049) + Penicillin/Streptomycin (Thermo Fisher, P0781) + 0.5x N2 supplement (Thermo Fisher, 17502048) + 1x B27 (Thermo Fisher, 12587010) + 1x Glutamax (Thermo Fisher, 35050061) + 1x MEM NEAA (Thermo Fisher, 11140035) + 2.5ug/ml Insulin (Millipore, I9278)] and 1% matrigel. SLNs were cultured on an orbital shaker with a rotation orbit diameter of 25mm set at a speed of 64 rpm. At later time points the medium was switched to IDM+A medium (IDM-A medium containing 1x B27 with vitamin A (Thermo Fisher, 17504044) instead of 1x B27 without vitamin A (Thermo Fisher, 12587010)). Medium was replaced every second day till day 20. From day 20, the medium was changed twice a week.

### SLN culture in different plate formats

SLNs were generated using various commercially available microwell plates. These included Elplasia plates (Corning, 4441), Aggrewell plates (Stemcell Technologies, 34411 and 34415) and PEG microwell plates (Sunbiosciences or Insphero, Gri3D® hydrogel microcavity plates) with microwell diameters of 300 μm, 500 μm, 800 μm, 1000 μm, and 1500 μm, as well as 96-well U-bottom plates coated with polyethylene glycol (PEG) (Sunbiosciences or Insphero hydrogel custom plates) and ultra-low attachment (ULA) 96-well U-bottom plates (SBio, MS-9096UZ). For each plate format, the cell suspension density was adjusted to ensure a seeding density of 10-12 cells per microwell. Culture plates were handled according to the respective manufacturer’s instructions. Subsequently, all cultures underwent the standard SLN differentiation protocol as previously described, adjusted for medium volume as per manufacturer’s instructions. SLNs were live imaged on day 6 and day 8 on Nikon Eclipse Ti inverted microscope.

### Mouse SLN generation

The mouse embryonic stem cell line CGR8 (Sigma, 07032901) was maintained under feeder-free conditions in 2i LIF medium. This medium consisted of high-glucose DMEM (Gibco, 61965026) supplemented with 1% non-essential amino acids (Thermo Fisher, 11140035), 1% sodium pyruvate (Thermo Fisher, 11360039), 0.1 mM β-mercaptoethanol (Thermo Fisher, 31350010), 10% fetal bovine serum (FBS) (Thermo Fisher, 16141079), 1,000 U/ml LIF (Sigma, ESG1106), 3 μM CHIR99021 (Millipore, 361571), and 1 μM PD0325901 (Selleck Chemicals, S1036). Mouse ES cells were passaged every 2 days using Accutase (Gibco, cat. no. 12605010) for dissociation and seeded at a density of 30,000-40,000 cells per 6-well plate in MEF-conditioned medium supplemented with 2i LIF. ES cells were routinely tested for mycoplasma contamination. For SLN differentiation, 5-8 cells were seeded per microwell in 500 μm microwell plates (Sunbiosciences or Insphero, Gri3D® hydrogel microcavity plates) in NIM supplemented with CEPT. Two days post-seeding, the medium was changed to NIM. The medium was refreshed every other day.

### Differentiation of SLNs into brain organoids Forebrain organoids

SLNs were generated following the protocol previously described. On day 6, the SLNs were treated with NIM supplemented with 10 μM SB 431542 (Tocris, 1614), 3 μM IWR1 (Sigma-Aldrich, 681669) and 5% Matrigel. On day 8, 500-600 SLNs were transferred from microwells to an ultra-low attachment 9 cm dish in IDM-A supplemented with 10 μM SB 431542, 3 μM IWR1, and 1% Cultrex (R&D systems, 3533-005-02). The medium was refreshed every other day, and cultures were maintained on an orbital shaker with a rotation orbit diameter of 25mm set at a speed of 64 rpm. From day 16, the medium was switched to 1 μM CHIR (Tocris, 4423) in IDM-A supplemented with 1% Cultrex and the medium was changed every other day. From day 18 onwards, the medium was switched to IDM-A supplemented with 1% Cultrex and the medium was changed every other day, From day 30 onwards, the medium was switched to IDM+A, supplemented with 1% Cultrex, and the medium was refreshed twice a week. Small retinal organoids formed in the forebrain organoid cultures and were removed based on morphology.

### Midbrain organoids

SLNs were generated as previously described. On day 8, 250-300 SLNs were transferred to an ultra-low attachment 9 cm dish. The culture medium was switched from NIM to IDM-A, supplemented with 1.5 μM CHIR (Tocris, 4423), 0.5 μM SAG (Stem Cell Technologies, 73412), 100 ng/ml FGF8b (Peprotech, 100-25) and 1% matrigel. The medium was refreshed every other day, and cultures were maintained on an orbital shaker with a rotation orbit diameter of 25mm set at a speed of 64 rpm. The medium, supplemented with growth factors and extracellular matrix, was refreshed every other day until day 16. From day 16 onwards, to test the effect of long term presence of soluble ECM in the medium, the cultures were maintained with or without 1% Matrigel till day 60. Starting on day 16, the medium was switched to IDM+A and replaced twice a week. From day 30 onwards, the medium was refreshed twice a week.

### Hindbrain organoids

SLNs were generated as previously described. On day 6, the SLNs were treated with NIM supplemented with 10 μM SB43, 2.5 μM Dorsomorphin (Sigma-Aldrich, P5499), 1.5 μM CHIR, 7 μg/ml insulin (Sigma-Aldrich, I9278), 20ng/mL FGF2 (Peprotech, 100-18B), 20ng/mL EGF (R&D, 236-EG-200) and 5% Matrigel. On day 8, 500-600 SLNs were transferred to an ultra-low attachment 9 cm dish containing IDM-A supplemented with 10 μM SB43, 2.5 μM Dorsomorphin, 1.5 μM CHIR, 7 μg/ml insulin, 20ng/mL FGF2, 20ng/mL EGF and 100 ng/ml FGF8b (Peprotech, 100-25). The medium was refreshed every other day, and cultures were maintained on an orbital shaker with a rotation orbit diameter of 25mm set at a speed of 64 rpm. Beginning on day 14, the medium was switched to 10 μM SB43, 2.5 μM Dorsomorphin, 1.5 μM CHIR, 7 μg/ml insulin, 20ng/mL FGF2 and 20ng/mL EGF and 100 ng/ml FGF19. The medium was refreshed every other day, and cultures were maintained on an orbital shaker. On day 22, the medium was switched to IDM-A and changed every other day until day 30. From day 30 to day 60, IDM-A was supplemented with 100 ng/ml SDF1a (Peprotech, 300-28A) and 0.5 ng/ml T3 (Merck, T-074-1ML) was refreshed two times a week. From day 60 onwards, the medium was switched to IDM-A.

### Retinal organoids

SLNs were generated as previously described. From day 6 to day 12, the SLNs were treated with dual SMAD inhibition treatment with 2.5 μM Dorsomorphin (Sigma-Aldrich, P5499), 10 μM SB 431542 (Tocris, 1614) and 3 μM IWR1 (Sigma-Aldrich, 681669) in NIM. On day 8, 500-600 SLNs were transferred to a 9 cm ultra-low attachment dish and cultured on an orbital shaker with a rotation orbit diameter of 25mm set at a speed of 64 rpm. Beginning on day 12, the medium was switched to IDM-A supplemented with 50 ng/ml BMP4 (Peprotech, 120-05ET), and it was replaced every other day. On day 24, the medium was switched to IDM-A and refreshed every other day.

For long term maturation, the medium was transitioned to a series of differentiation media based on previously published protocols^39,40,54^. Specifically, from day 40, cultures were maintained in retina differentiation medium 1 [RDM1 - DMEM high glucose, GlutaMAX, F-12 Nutrient Mixture (Ham), 1x B27 without vitamin A, 1x NEAA Solution, 10% ES FBS (Thermo Fisher, 16141079), 1x Sodium Pyruvate 100 mM, 1x Penicillin/Streptomycin, 1x Amphotericin B, freshly supplemented with 100 mM Taurine (Sigma, T8691-25G)] with medium changes every other day. To test the effect of ECM on organoid growth, the cultures were maintained with or without 1% Matrigel till day 60.

From week 10 onwards, the medium was switched to RDM2 (RDM1 supplemented with 1 μM Retinoic acid (Sigma-Aldrich, R2625) and 100mM Taurine). From week 14, organoids were transferred to an ultra-low attachment 24-well plate, with individual organoids in separate wells under static conditions to prevent fusion and mechanical stress, and the medium was switched to retina differentiation medium 3 [RDM3 - DMEM, high glucose, GlutaMAX, F-12 Nutrient Mixture (Ham), 1x N2 supplement, 1x NEAA Solution, 10% ES FBS, 1x Sodium Pyruvate 100 mM, 1x Penicillin/Streptomycin, 200mg/ml Albumax (Thermo Fisher, 11021029), Amphotericin B, freshly supplemented with 0.5 μM Retinoic acid and 100mM Taurine]. At this time point, organoids with a shiny neuroepithelial rim were selected under a stereo microscope to further culture. Organoids with dense morphology or small organoids with very small neuroepithelium perimeter were discarded.

### Morphogen treatment experiment

For each condition, 500-600 SLNs on day 8 were transferred to one 6 cm dish in IDM-A supplemented with 3% Matrigel and the following individual morphogens: 3 μM CHIR (Tocris, 4423), 0.5 μM SAG (Stem cell Technologies, 73412), 2.5 μM DAPT (Stem cell Technologies, 72082), 1 μM Retinoic acid (Sigma-Aldrich, R2625-50MG), 10 ng/ml Recombinant Human BMP-4 (Peprotech, 120-05ET), 100 ng/ml Recombinant Human/Murine FGF-8b (Peprotech, 100-25), 2.5 μM IWP2 (Selleckchem, S7085), 0.5 μM LDN193189 (Stem cell Technologies, 72147), 10 μM SB 431542 (Tocris, 1614), 20 ng/ml Recombinant Human FGF-2 (Peprotech, 100-18B), 50 ng/ml Activin A (Peprotech, 120-14E-100UG) and untreated. The medium was changed every other day, and SLNs were harvested on day 16 for single-cell RNA sequencing and immunostaining.

### Reaggregation experiments

To evaluate the reaggregation capacity of differentiated SLNs, 500-600 SLNs were harvested on day 10 of the standard differentiation protocol and dissociated into single cells using the Papain-based Neural Tissue Dissociation Kit (Miltenyi Biotec, 130-092-628) according to the manufacturer’s instructions. The resulting cell suspension was centrifuged at 300 g for 5 minutes, and the cell pellet was resuspended in NIM supplemented with CEPT. These cells were then seeded into 500 μm microwells at a density of 100 cells per microwell. The following day, the culture medium was replaced with NIM supplemented with 5% Matrigel. The medium was subsequently changed every other day. Reaggregating cells were imaged for SOX2-GFP expression and under brightfield illumination on day 6 post-seeding using Nikon Eclipse Ti inverted microscope.

### Live imaging for interkinetic nuclear migration assay

To visualize cell morphology and tissue organization, WTC-CAAX-RFP and WTC-ZO1-GFP iPSC lines were mixed at a ratio of 1:100 before seeding into the microwell plates. Following the standard SLN differentiation protocol, SLNs were carefully transferred to glass-bottomed plates (Cellvis, P96-1.5H-N) for live imaging. Prior to imaging, SLNs were stained with BioTracker 650 red nuclear dye (Sigma-Aldrich, SCT119) according to the manufacturer’s instructions. Live imaging was performed using a Leica Stellaris 8 microscope with 25x water immersion lens within a controlled incubation chamber maintained at 37 °C and 5% CO2.

### Immunocytochemistry

#### Whole mount staining

Organoids were collected from microwell plates in microfuge tubes, washed with 1X Phosphate Buffered Saline (PBS, Sigma-Aldrich, P5493) and fixed with 4% Paraformaldehyde (Electron Microscopy Sciences, 15710) for 2 hours at 4C. Samples were washed with PBS and stored in PBS supplemented with 0.2% Sodium Azide (Sigma- Aldrich, 71290) until used for immunostaining. Fixed samples were incubated in blocking solution [PBS supplemented with 3% donkey serum (Jackson ImmunoResearch, 017-000-121) and 0.1% Triton-X (Millipore Sigma, T8787)] for 1 hour at room temperature in microfuge tubes on a shaker. Samples were then incubated with primary antibodies diluted in blocking solution at 4°C overnight on a shaker. Next day, samples were washed 3 times with PBS for 5 minutes. Samples were then incubated with secondary antibodies and Hoechst 33342 (ThermoFisher Scientific, H3570) diluted in blocking solution at 4°C overnight on a shaker followed by 3 washes with PBS for 5 minutes each. Samples were stored in PBS until imaged. For clearing, PBS was removed and samples were submerged in Cubic mount solution^55^ [250g sucrose 50%w/v Sucrose, 25%w/v Urea, 25%w/v N,N,N′,N′-tetrakis(2-hydroxypropyl)ethylenediamine] overnight at room temperature. Samples were stored at room temperature until imaged. Samples were transferred to glass bottom 96-well plates (Cellvis, P96-1.5H-N) and imaged using Nikon Eclipse Ti inverted microscope.

#### Sections

Samples were washed with 1X Phosphate Buffered Saline (PBS, Sigma-Aldrich, P5493) and fixed with 4% Paraformaldehyde (Electron Microscopy Sciences, 15710) overnight at 4°C. Samples were washed with PBS and stored in PBS supplemented with 0.2% Sodium Azide (Sigma-Aldrich, 71290) until used for immunostaining. Fixed samples were embedded into histogel (Epredia, HD-4000-012) wells generated using custom made molds specific to the size of the organoid type^56^. Samples were dehydrated and paraffin-embedded using an automated tissue processor (Leica HistoCore PEARL) and an embedding station (Medite TES99). Paraffin blocks were sectioned at 5 μm on a microtome (Thermo Microm HM355S microtome). Sections were mounted on slides (Fisher scientific, 11976299) and dried.

For automated staining, sections were subjected to multiplexed immunofluorescence using Opal dyes (Akoya Biosciences) on a Roche Ventana Discovery Ultra stainer following manufacturer’s instructions. For multiplexed immunofluorescence, slides underwent deparaffinization, heat-induced antigen retrieval, blocking, antibody incubation, Opal dye application, and antibody neutralization/denaturation steps between each staining cycle. Slides were mounted with ProLong™ Gold Antifade Mountant (Invitrogen P36930) and dried prior to imaging. Slides were imaged using a Vectra Polaris (Akoya Biosciences Perkin Elmer) at 20x magnification for all 7 colors (Opal 480, Opal 520, Opal 570, Opal 620, Opal 690 and Opal 780). Laser exposure and intensity settings were adjusted on multiple slides per staining panel. Channels were unmixed and images tiled with PhenoChart (v1.0.12) and inForm (v2.4). Raw images were saved as .qptiff and fused in HALO (Indica labs, v3.6.4134.396). Image analysis of multiplexed immunofluorescence images was performed with HALO.

For manual staining, after antigen retrieval sections were incubated in blocking solution [PBS supplemented with 3% donkey serum (Jackson ImmunoResearch, 017-000-121) and 0.1% Triton-X] for 1 hour at room temperature. Samples were then incubated with primary antibodies diluted in blocking solution at 4°C overnight. The next day, samples were washed 3 times with PBS for 5 minutes. Samples were then incubated with secondary antibodies and Hoechst 33342 (ThermoFisher Scientific, H3570) diluted in blocking solution for 1 hour at room temperature followed by 3 washes with PBS for 5 minutes each. Slides were mounted with ProLong™ Gold Antifade Mountant (Invitrogen P36930) and were stored at 4°C until imaged. Samples were imaged using Nikon Eclipse Ti inverted microscope or Olympus Slideview VS200 Universal Whole Slide Imaging Scanner. Image analysis was performed with HALO (Indica labs, v3.6.4134.396).

A list of primary and secondary antibodies is provided in Supplementary table 1.

### Image analysis

Live imaging of the single lumen organoids was performed using the Nikon Eclipse Ti inverted microscope with 10x magnification equipped with Hamamatsu ORCA-Fusion C14440-20UP camera and an incubation chamber at 37 °C, 5% CO2. Cells were imaged on day 4, 6, 8 and 10 of differentiation. Tiles images of each well were analyzed with ImageJ 2.9.0 using built-in plugins. To determine organoids’ size over time, ten brightfield images were thresholded and segmented. The distance between the two parallel planes restricting the organoid, namely the Feret’s diameter, was measured on the detected particles. Similarly, lumen’s diameter was measured using the ZO1-GFP reporter signal. Data was plotted in Python (v3.11.4) using Matplotlib (v3.8.3) and Seaborn (v0.13.2) within a Jupyter Notebook.

Quantification of neuroepithelium classes [no neuroepithelium (E), SLN (S) or multiple lumina (M)] was performed using Aivia imaging software (v14.1.9, Leica Microsystems) on brightfield images acquired with a Nikon Eclipse Ti inverted microscope. Initially, pixel classifiers were trained on a representative image from each well size through regional annotation of background, well features and SLNs, and then applied to the acquired image set for each well size. Next, the obtained channels were input in Aivia’s built-in “Cell count” recipe with optimized detection parameters to generate a set of quantifiable objects with minimal background signal. Finally, extracted objects from one image per well size were used as an input to train object classifiers capable of distinguishing and quantifying single- and multilumen neuroepithelium.

These trained object classifiers were applied on the corresponding well size images datasets, with class allocation iteratively refined to ensure classification accuracy. Data for microwell size was plotted in RStudio (v2024.12.0) using dplyr (v1.1.4) and ggplot2(v3.5.2). Data from different iPSC lines was plotted in Python (v3.11.4) using Matplotlib (v3.8.3) and Seaborn (v0.13.2) within a Jupyter Notebook.

For regional organoids, marker quantification analysis was done using the HALO software v3.6.413. A classifier was trained for each replicate and timepoint using the Random Forest classifier, by drawing sample areas across organoid sections in order to detect organoids, their necrotic core and background glass. Necrotic cores were only separated for d60 samples as d20 samples did not show any signs of a deteriorating core. Nuclear and marker detection was run on the organoid classified area excluding necrotic cores and background using the CytoNuclear FL module v.2.0.12 for nuclear marker as well as Area Quantification FL c2.3.4 for cytosolic marker. For the CytoNuclear FL analysis of nuclear markers, nuclei were detected using the DAPI stain. Minimum nuclei size for detection has been set to 9 µm², a nuclear contrast threshold of 0.525 was applied and nuclear segmentation was applied. Minimum detection intensities were selected for each marker with the help of the Real-time Tuning feature to resemble the actual signal. Due to varying signal intensities across timepoints minimum detection intensities for markers had to be adjusted slightly to better match the sample’s fluorescent signal. For cytoplasmic marker quantification, no cell detection was performed on the organoid section of the classifier. As before minimum detection intensities were set according to the apparent signal and a blur radius of 0.25 was applied. Both analyses were performed on individual organoids using the TMA core function. The data was exported, analyzed and visualized using R (version 4.4.1 (2024-06-14)).

For measuring retinal organoid size, organoid brightfield images were segmented using Cellpose v4.0.6 with the SAM-based CellPose Model. Images were downsampled 4× and quantile-normalized before segmentation, with an initial diameter of 120 pixels, adapted in a second pass for the day-specific median diameter for adaptive segmentation. Segmentation masks were post-processed to remove border-touching objects, and organoid diameters were computed from the masks and corrected for downsampling.

For quantifying the nuclear layer in retinal organoids, NRL and SOX9 location analysis was done using the HALO software v3.6.413. A classifier was trained for each replicate image using the Random Forest classifier, by drawing sample areas across organoid sections in order to separate organoids from background glass. The classifier was set up to generate an annotation around the perimeter of the organoid section. Separate analyses were created for SOX9 and NRL for each replicate using the CytoNuclear FL module v.2.0.12. Cell detection parameters were set up to identify nuclear signals. Using the “Real-time Tuning” feature, values had to be adjusted slightly across replicates depending on the fluorescent intensities to accurately represent the apparent fluorescent signal. Before running the analysis, images were divided into smaller sections using the “Segment TMA (tissue mold array)” function. Subsequently, the analysis was performed on the individual TMA cores. The initial analysis was followed up by an infiltration analysis. The object data was exported and analyzed using R (v4.4.1). Briefly, based on the TMA core location unique Organoid IDs were generated and assigned to the respective data. The data was cleaned up by removing N/A values, duplicate entries and applying organoid specific exclusions as well as reintroducing 0-value organoids. Exclusions were applied in cases of missing data for either of the markers. 0-value organoids were added for organoids that went through analysis but had no marker-positive cells and thus did not appear in the exported data. Cell counts were divided into 1 µm bands and plotted as a heatmap.

### Single cell RNA sequencing

For analyzing the day 10 SLNs, cells were differentiated as described above. On day 10, 250-300 SLNs were transferred to a microfuge tube and washed with PBS (Gibco, 14190) and dissociated into single cells using Papain-based neural tissue dissociation kit (Miltenyi Biotec, 130-092-628). Dissociated cells were run through 30μm cell strainer (CellTrics, 04-0042-2316), spun at 300g for 5 minutes and resuspended in PBS + 0.1% bovine serum albumin (Gibco, 15260-037). Cell density was adjusted to 1 million cells/ml and 5000 cells were captured for RNA sequencing using 10X Genomics Single Cell 3’ v3.1 protocols with feature barcode technology.

For single cell analysis of the SLNs after morphogen treatment, half the cultured SLNs (more than 100) were transferred to a microfuge tube, washed with PBS, and dissociated into single cells using a Papain-based neural tissue dissociation kit. Dissociated cells were run through a 30-μm cell strainer, centrifuged at 300g for 5 minutes, and resuspended in PBS + 0.05% bovine serum albumin. Samples were counted, pooled, and barcoded using the 3’ CellPlex Kit (10X Genomics, 1000261) according to the manufacturer’s protocol. Briefly, samples were washed with 10% FBS + PBS solution, centrifuged at 300g for 5 min at 4°C. Individual samples were resuspended in Cell Multiplexing Oligo, incubated at room temperature for 5 minutes, washed with 10% FBS + PBS, and centrifuged at 300g for 5 min at 4°C. Samples were resuspended, counted, and pooled in equal ratios for encapsulation using 10X Genomics Single Cell 3’ v3.1 protocols with feature barcode technology.

For single cell analysis of regionalised organoids, 3 organoids were pooled, washed with PBS, and dissociated into single cells using a Papain-based neural tissue dissociation kit. The time for dissociation was optimised for the organoids type. Dissociated cells were run through a 30-μm cell strainer, centrifuged at 300g for 5 minutes, and resuspended in PBS + 0.1% bovine serum albumin.Cell density was adjusted to 1 million cells/ml and 10000-20000 cells were captured for RNA sequencing using 10X Genomics Single Cell 3’ v4 protocols with feature barcode technology.

For all experiments, libraries were prepared according to manufacturer’s recommendations (10X Genomics Single Cell 3’ v3.1 or v4). Samples were sequenced on an Illumina NovaSeq6000 or NovaSeqX plus. Sequence FASTQ files were processed using Cellranger. Single cell transcriptomes were mapped to human (GRCh38/hg19) reference transcriptomes.

### Single cell data analysis

For the day-10 SLN scRNA-seq data, the analysis was mostly done with Seurat (v5) in R. The quality control (QC) metrics including detected transcript numbers (nUMIs), detected gene numbers (nGenes), mitochondrial transcript proportion (Mito%) and ribosomal protein transcript proportion (RP%) were firstly calculated per cell. QC filtering on cells was then performed by requiring nGenes between 2500 and 9500, as well as Mito% lower than 15%. Gene counts were normalized by the total transcript number per cell, followed by log-transformation. Cell cycle phase scores were calculated using the *CellCycleScoring*() function in Seurat, given genes annotated to be G2M and S phases related (*cc.genes.updated.2019* in Seurat). Highly variable genes (HVGs) were identified and the top-3000 HVGs were selected. The log-normalized expression values were scaled per gene across cells with the *ScaleData()* function. Principal Component Analysis (PCA) was applied to the scaled expression of HVGs, and the first 20 principal components (PCs) were chosen to perform uniform manifold approximation and projection (UMAP) to generate the 2D embedding for visualization.

For the scRNA-seq data of the day-16 SLN with different signaling treatment, the preprocessing and analysis was mostly done with Seurat (v5) in R. First, similar QC metrics were calculated as described above. QC filtering on cells was performed by requiring nGenes between 3000 and 10000, as well as Mito% lower than 10%. The gene counts were used as the input to project to the previously established SCANVI model (https://zenodo.org/records/15004818)^28^ of the first-trimester developing human brain atlas^27^ with scArches^57^, to derive the projected SCANVI latent representations of the data. The kNN were constructed and used to generate UMAP embedding for visualization and perform Leiden clustering (resolution=1). The resulting clusters were annotated based on canonical marker expression and the transferred cell class and regional labels from the developing human brain atlas. The label transfer was based on the bipartite weighted k-nearest neighborhood (kNN) between SLN data and the primary reference as implemented in HNOCA-tools28. The same projected SCANVI latent representation was also used to estimate the anterior-posterior scores as well as the dorsal-ventral scores, both of which are based on an elastic net model^51^ trained using the radial glia of the first-trimester developing human brain atlas^27^.

For the scRNA-seq data sets of the regionalized SLNs, the preprocessing and analysis was mostly done with Scanpy in Python. First, the four individual data sets of regionalized SLNs were concatenated with the outer manner. Similar QC metrics were calculated as described above. QC filtering on cells was performed by requiring nGenes between 1500 and 8000, as well as Mito% lower than 15%. Gene counts were normalized by the total transcript number per cell (*target_sum*=1e4), followed by log-transformation. Highly variable genes (HVGs) were identified using the Seurat v3 algorithm (*flavor*=’seurat_v3’) and the top-4000 HVGs were selected. Among them, the 2755 HVGs detected in all the datasets (common HVGs) were used for the upcoming analysis. PCA was applied to the log-normalized expression of the common HVGs. The top-20 PCs were used to construct kNN, which was used to perform UMAP embedding for visualization, and leiden clustering (resolution=0.5). The resulting clusters were annotated based on canonical marker expression and the transferred cell class and regional labels from the developing human brain atlas. The label transfer was done in a similar procedure as above.

To characterize the regionalized SLNs scRNA-seq datasets, we adapted the established procedure in HNOCA by comparing the datasets to the first-trimester developing human brain cell atlas. In brief, for each of the four regionalized SLN datasets, a normalized presence score was calculated for each cell in the primary atlas, based on the projected SCANVI latent representation and the corresponding bipartite kNN with the primary atlas, as described previously and implemented in HNOCA-tools^28^. The presence scores quantify how well a cell state represented by each reference cell presents in each of the regionalized SLN dataset. The resulting scores were then summarized into mean presence scores per cell cluster. Next, the cluster-level presence score profile per regionalized SLN dataset was clustered together with the HNOCA datasets^28^. To quantify transcriptomic similarity to the primary reference for each cell in the regionalized SLN datasets, a matched primary metacell transcriptomic profile was generated for each cell in the SLN scRNA-seq datasets, as implemented in HNOCA-tools^28^.

Pearson correlation coefficient was calculated for each cell, between its actual measured transcriptomic profile and its matched primary transcriptomic profile across highly variable genes in the first-trimester developing human brain atlas. The same procedure was applied to the HNOCA data^28^. To correct for the confounding effect of detected gene number per cell on the calculated correlation, a linear model was fitted with the calculated Pearson correlation as the dependent variable and log-transformed detected gene number as the independent variable, given the concatenated data of the regionalized SLNs scRNA-seq data and the HNOCA data.

### Residuals of the model were extracted as the corrected correlation

For the retinal organoid scRNA-seq data, the preprocessing and analysis was mostly done with Seurat in R. First, similar QC metrics were calculated as described above. QC filtering on cells was performed by requiring nGenes between 1500 and 9000, nUMIs below 40000, as well as Mito% lower than 10%. After log-normalizing the gene counts, the top-5000 HVGs were identified. PCA was applied to the scaled expression of the HVGs, with the top-20 PCs selected to perform leiden clustering (resolution=0.5) and generate the UMAP embedding for visualization. The resulting clusters were annotated based on canonical marker expressions. To compare with other retinal organoid protocols, we projected the newly generated scRNA-seq data to the previously published retinal organoid time course cell atlas^45^, based on the CSS representation model^28^ of the published atlas. With the bipartite weighted kNN based label transfer as described above but based on the projected CSS representation, cell type labels were transferred from the published retinal organoid atlas to the newly generated retinal organoid data. Next, the published atlas was split into eight partitions based on the rounded organoid ages in months. For each of the partitions, a bipartite weighted kNN was built between the newly generated data and the partition, so that a matched transcriptomic profile was generated for each cell in the newly generated dataset using the same procedure as described above. For each cell in the newly generated dataset, a Pearson correlation coefficient was calculated between its observed transcriptomic profile and the matched transcriptomic profile per reference partition, across HVGs defined in the published atlas. The distributions of the resulting transcriptomic similarities were used to characterize matching maturity of the newly generated data in the time course of the published atlas.

### Statistical Analysis

Statistical parameters such as the value of n, mean, standard deviation (SD), p values, and the statistical tests used are reported in the figure legends. For hiPSC differentiation experiments, the “n” refers to the number of independent hiPSC differentiation experiments analyzed.

### Supplementary Tables

1. Supplementary table 1 - List of antibodies used in this study

2. Supplementary table 2 – List of HNOCA datasets

## Acknowledgements

We thank 360 Genomics labs, Genetics and Genomics core and Marina Bellavista at Roche Pharma Research and Early Development (pRED) for their help with single cell RNA sequencing. We thank all lab members for helpful discussion, feedback and resource sharing. To refine the textual clarity of early drafts, some contributors used generative AI tools, including Gemini and ChatGPT. Following the use of these resources, the resulting output was thoroughly revised by the authors to confirm the accuracy.

## Author contribution

J.R. designed and conducted hiPSC differentiation experiments, acquired data and performed data analysis. S.L. designed, performed and analyzed all regionalized organoid differentiations with J.R.. A.B. designed and performed experiments for forebrain and retina organoid protocol optimization. J.L. helped J.R. with optimization of hindbrain differentiation experiments. M.C. performed microwell size experiments and helped with image analysis. Y.J. performed live imaging on SLNs and helped with data analysis. I.C. performed multiplex immunohistochemistry experiments. J.G.C. provided input on the project. Z.H. and B.T. performed single cell RNA ssequencing analysis. J.R. and M.P.L. conceptualized the work, analyzed data and wrote the manuscript with input from all authors.

---

**Supplementary Figure 1.**
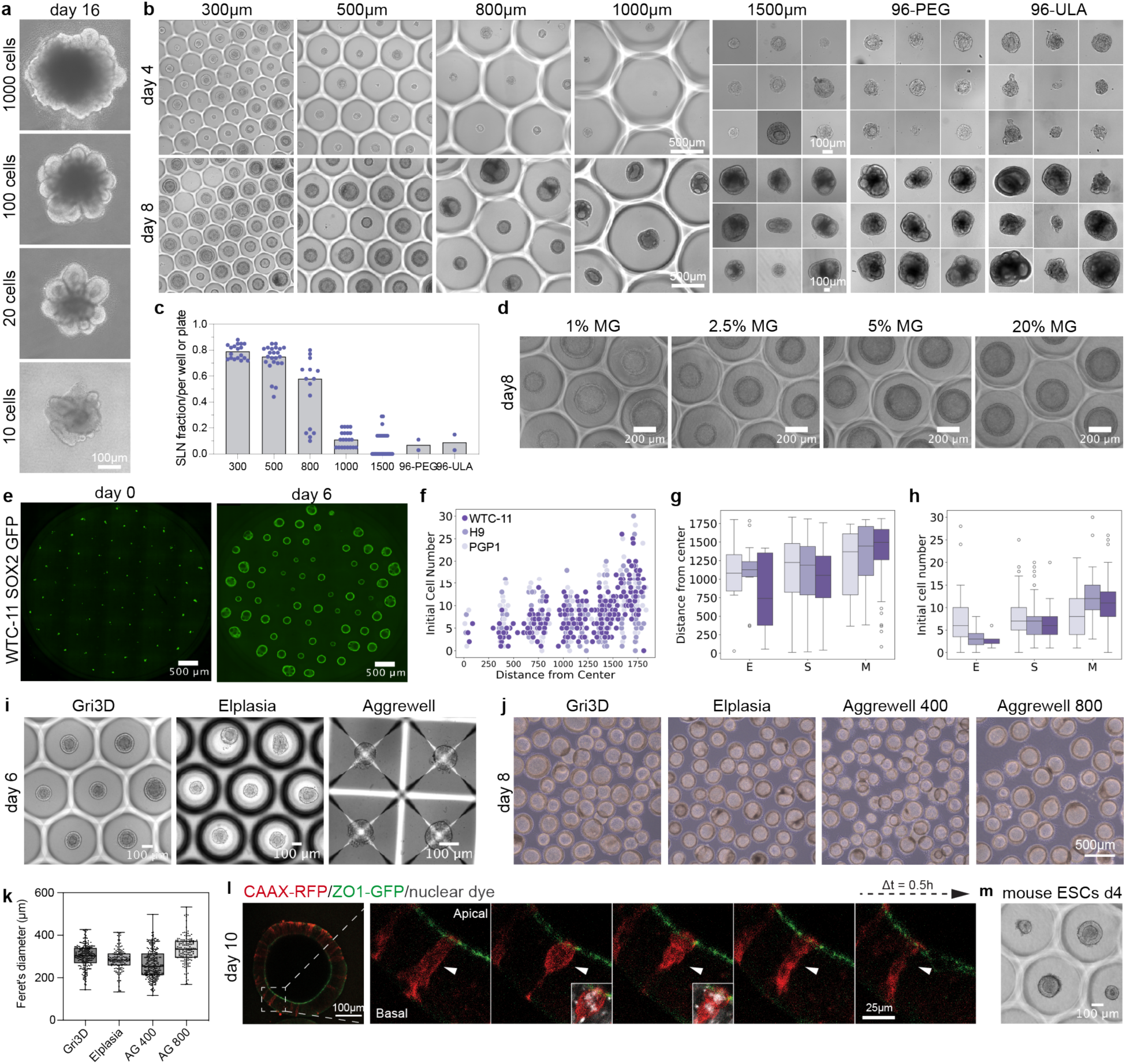
Generation of single-lumen neuroepithelium from human pluripotent stem cells. **(a)** Representative brightfield images illustrating the morphology of brain organoids derived using an unguided protocol (Lancaster et al. 2013) at day 16, demonstrating the effect of varying initial seeding cell number. **(b)** Representative brightfield images showing the formation of SLNs in microwells of different diameter sizes, in U-bottom PEG-coated 96 well plate and plastic ultra-low attachment (ULA) 96 well plates. **(c)** Quantification of SLNs formed in microwells of different sizes. n = 2-3 independent experiments, data points represent quantification of microwell plate wells or 96 well plates **(d)** Brightfield imaging of SLNs on day 8 of differentiation cultured in varying concentrations of Matrigel. **(e)** Representative fluorescence images showing WTC-11 SOX2-GFP-expressing cells seeded in a microwell on day 0 and their growth pattern at day 6 of differentiation. **(f)** Scatter plot showing the distribution of cell number settling into microwells in the well plate on day 0 for three cell lines. **(g)** Boxplot showing the relationship between the distance of neuroepithelium from the center of the well and the neuroepithelium category [no neuroepithelium (E), SLN (S) or multiple lumina (M)] in three cell lines. **(h)** Boxplot showing the distribution of initial seeding cell numbers in microwells that formed no neuroepithelium (E), SLN (S) or multiple lumina (M) on day 6 in three cell lines. **(i)** Representative brightfield images of SLN formation in PEG microwell plates (Gri3D, Sunbioscience or Insphero), ultra-low attachment microwells (Elplasia, Corning and AggreWell, StemCell Technologies) cell culture plate formats on day 6. **(j)** Representative brightfield images of transferred SLNs from different culture plate formats on day 8. **(k)** Quantification of the SLN size cultured in different plate formats (n≥130 neuroepithelium per condition). **(l)** Live imaging snapshots of SLNs derived from a mix of ZO1-GFP WTC-11 and CAAX-RFP WTC-11 iPSCs. Arrow indicates the dividing cells at the apical surface and exhibiting interkinetic nuclear migration. **(m)** Representative brightfield image of SLNs generated from mouse embryonic stem cells at day 4.

**Supplementary Figure 2.**
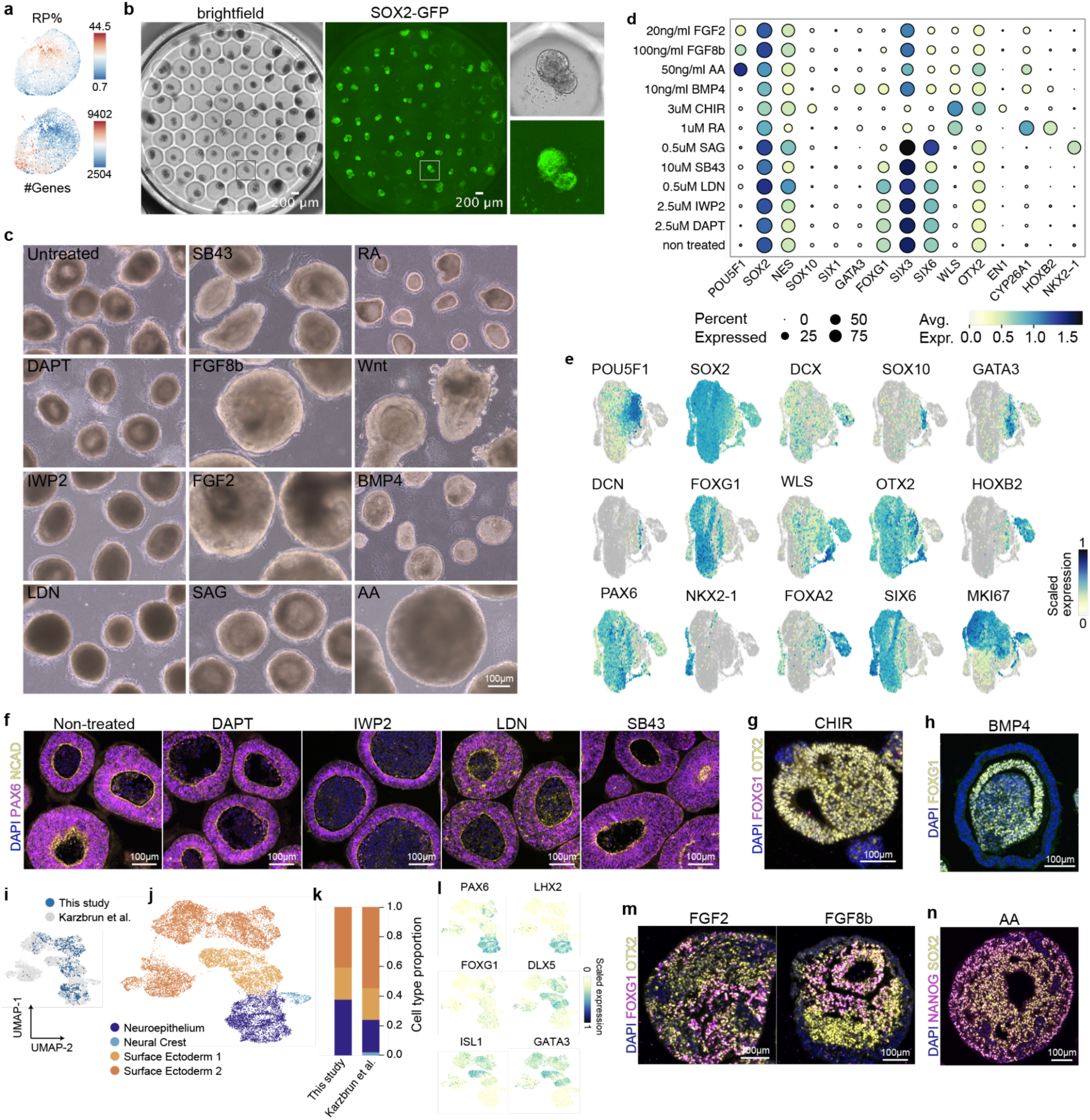
SLNs adopt a default anterior state but retain regional plasticity. **(a)** Feature plots overlaid on a UMAP embedding of single cells from day 10 SLNs, displaying the distribution of the percentage of ribosomal proteins (RP%) and the number of detected genes (#Genes) per cell, indicating cell quality. **(b)** Brightfield and fluorescence images showing SOX2-GFP-positive cells seeded in microwells following dissociation of day 10 SLNs. The inset displays the reorganisation of dissociated cells within a microwell into cystic neuroepithelial structures that retain SOX2-GFP expression. **(c)** Brightfield images illustrating the morphology of SLNs after 7 days of differentiation under various morphogen treatment conditions. **(d)** Dot plot showing the expression levels and percentage of expressing cells for selected regional identity marker genes across the different morphogen treatment groups. SB43 - SB431542, LDN - LDN193189CHIR - CHIR99021, RA - retinoic acid, AA - Activin A. **(e)** UMAP plots highlighting the expression of selected genes. **(f)** Representative immunofluorescence images showing the morphological changes and expression of PAX6 and N cadherin. Nuclei are counterstained with DAPI. **(g)** Immunofluorescence images showing FOXG1 and OTX2 expression after treatment with CHIR. **(h)** BMP4 treated SLNs stained with FOXG1 antibody. **(i)** UMAP projection showing integration of cells from this study with dataset from Karzbrun et al.^20^. **(j)** UMAP plot displaying cell type annotation from Karzbrun et al.. **(k)** Bar chart comparing the proportion of each cell type between this study and the reference dataset. **(l)** UMAP plots highlighting the expression of selected marker genes for identified cell types. **(m)** Immunofluorescence images showing FOXG1 and OTX2 expression after treatment with FGF2 and FGF8b. **(n)** Immunofluorescence images of SLNs stained for NANOG and SOX2 after Activin A treatment.

**Supplementary Figure 3.**
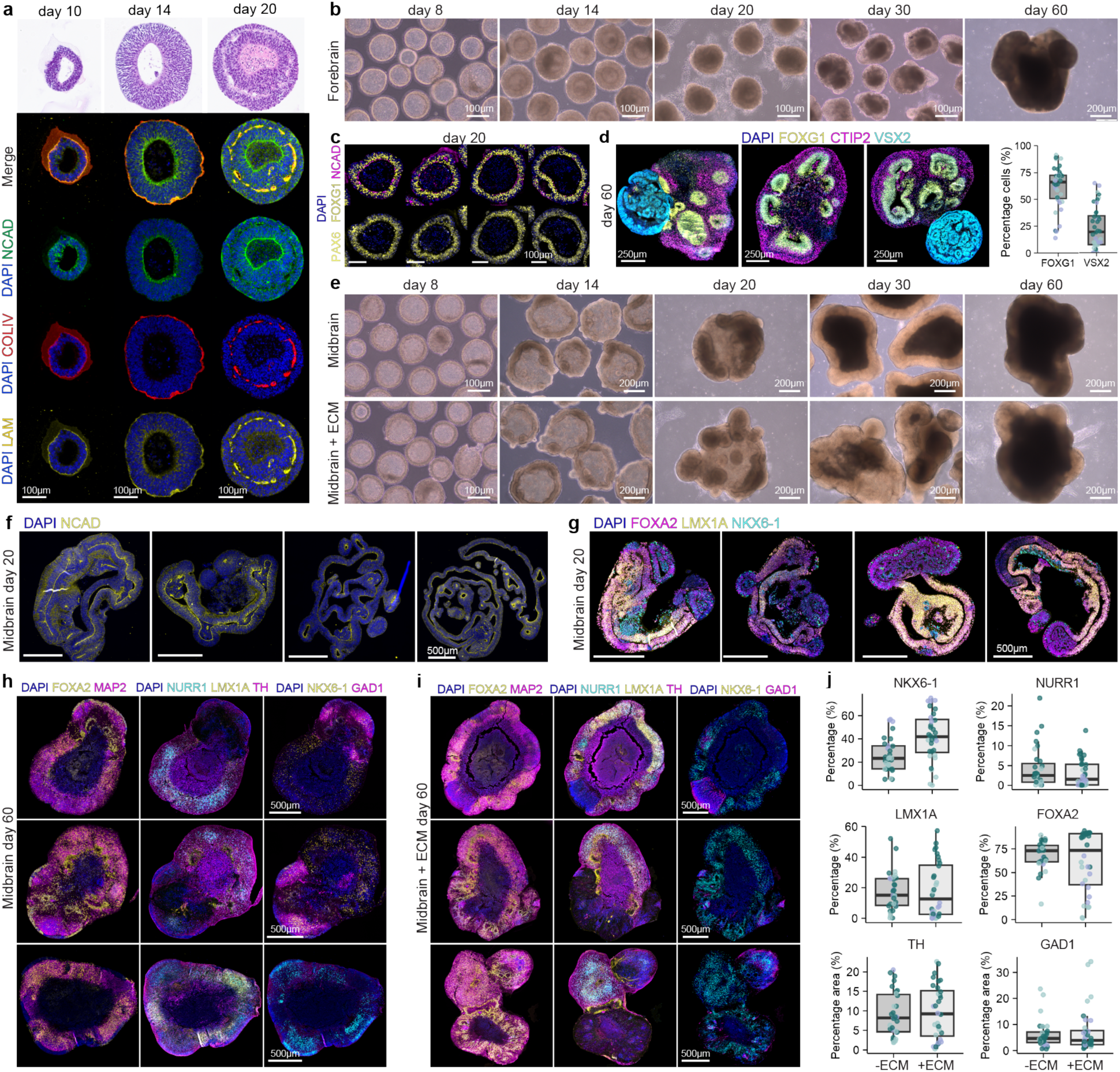
Generation of regionally specified brain organoids from SLNs. **(a)** Hematoxylin and Eosin (H&E) staining and Immunofluorescence analysis of SLNs showing the expression of markers apical polarity (NCAD), basement membrane (COLIV, Laminin), and nuclei (DAPI). Dissolved Matrigel was removed on day 10. **(b)** Brightfield images showing the morphological development of forebrain organoids over time. **(c)** Immunofluorescence analysis of day 20 forebrain organoids showing NCAD, PAX6 and dorsal forebrain marker FOXG1. **(d)** Immunofluorescence analysis and quantification for markers of dorsal forebrain (FOXG1) and retinal progenitor cells (VSX2). n= 3 independent experiments, 40 organoids. **(e)** Brightfield images showing the morphological development of midbrain organoids from SLNs over time. Both conditions were cultured in 1% Matrigel till day 16. **(f)** Representative immunofluorescence images of day 20 midbrain organoids stained for neuroepithelial marker SOX2 and apical marker NCAD. **(g)** Representative immunofluorescence images of day 20 midbrain organoids stained for FOXA2, LMX1A and NKX6-1. **(h)** Representative immunofluorescence images of day 60 midbrain organoids cultured long term without matrigel stained for midbrain markers. **(i)** Representative immunofluorescence images of day 60 midbrain organoids cultured long term with Matrigel stained for midbrain markers. **(j)** Quantification of midbrain markers on day 60 (30-35 organoids from 3 independent replicates).

**Supplementary Figure 4.**
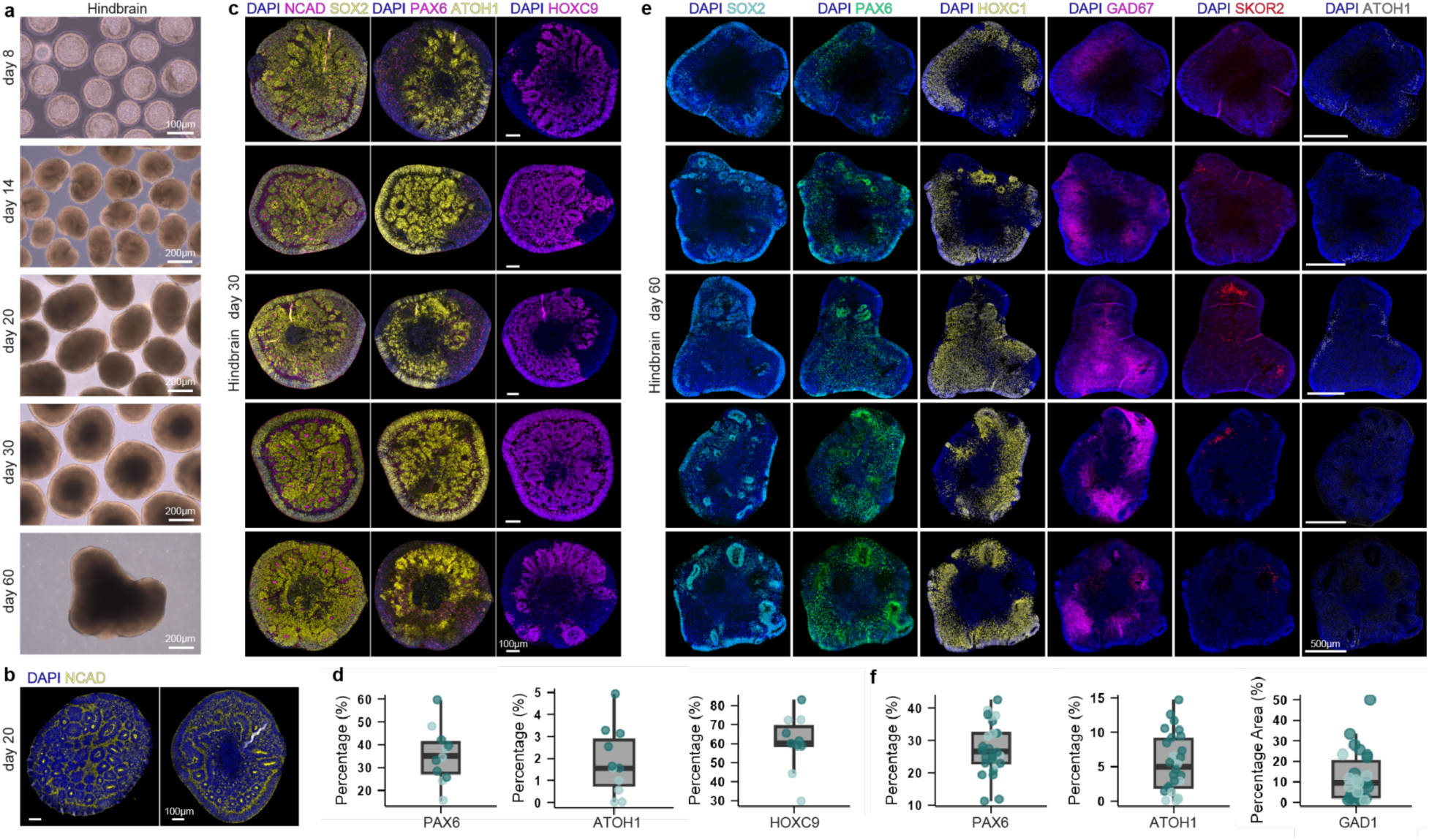
Generation of hindbrain organoids from SLNs. **(a)** Brightfield images showing the morphological development of hindbrain organoids from SLNs over time. **(b)** Immunofluorescence image showing expression of NCAD and SOX2 on day 20 of differentiation. **(c)** Representative immunofluorescence images of day 30 hindbrain organoids stained for PAX6, ATOH1 and HOXC9. **(d)** Quantification of hindbrain markers on day 20. n = 2 independent replicates). **(e)** Representative immunofluorescence images of day 60 midbrain organoids stained for hindbrain markers. **(f)** Quantification of hindbrain markers on day 60. n = 2 independent replicates).

**Supplementary Figure 5.**
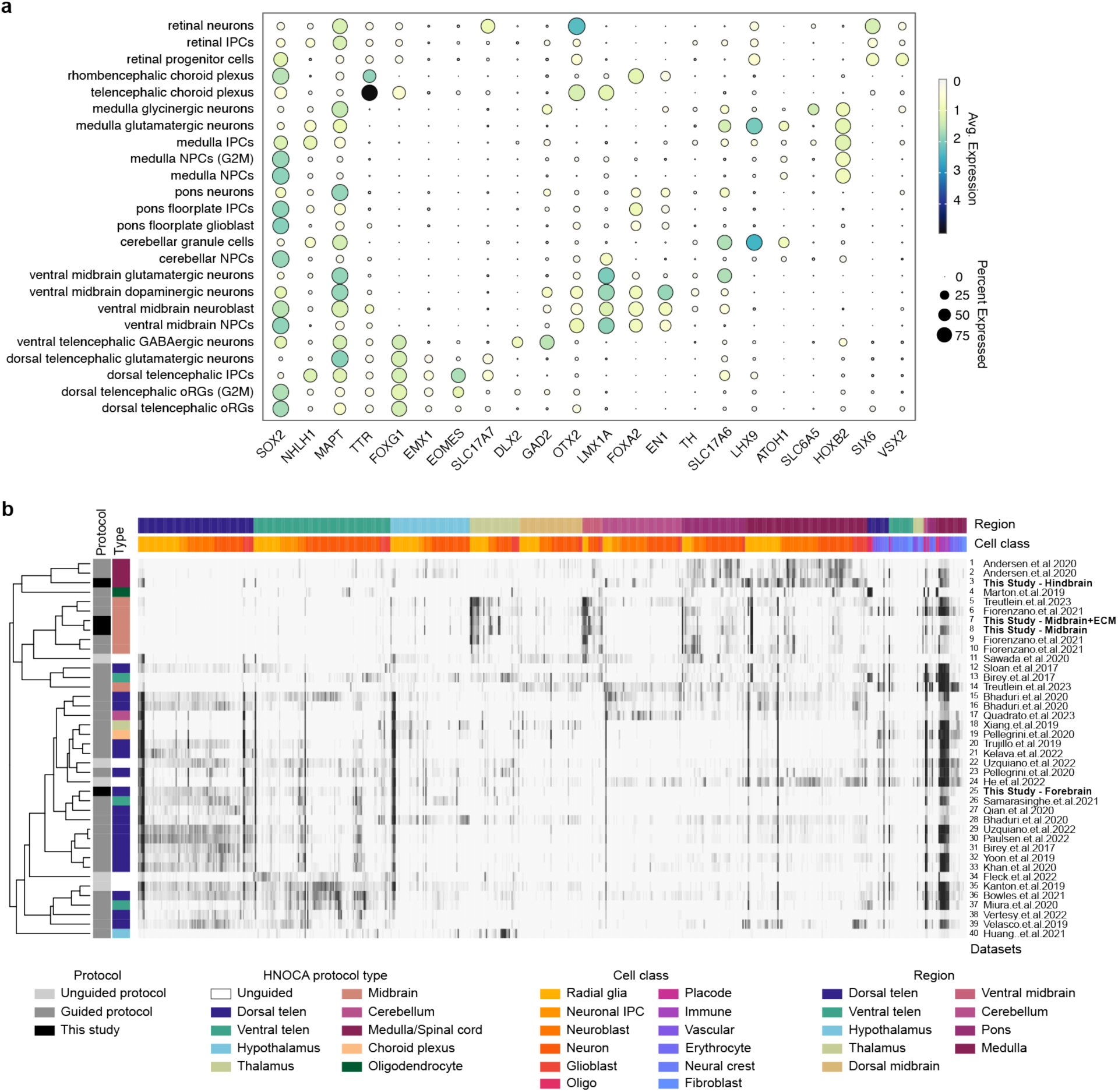
Single cell analysis of the regionally specified brain organoids from SLNs. **(a)** Dotplot showing gene expression levels of selected cell type markers across different cell types. **(b)** Clustering of HNOCA datasets (rows) with data generated in this study (rows) based on average presence scores (columns) of clusters in the primary reference. The heatmap shows average presence scores per cluster in the primary reference (columns).

**Supplementary Figure 6.**
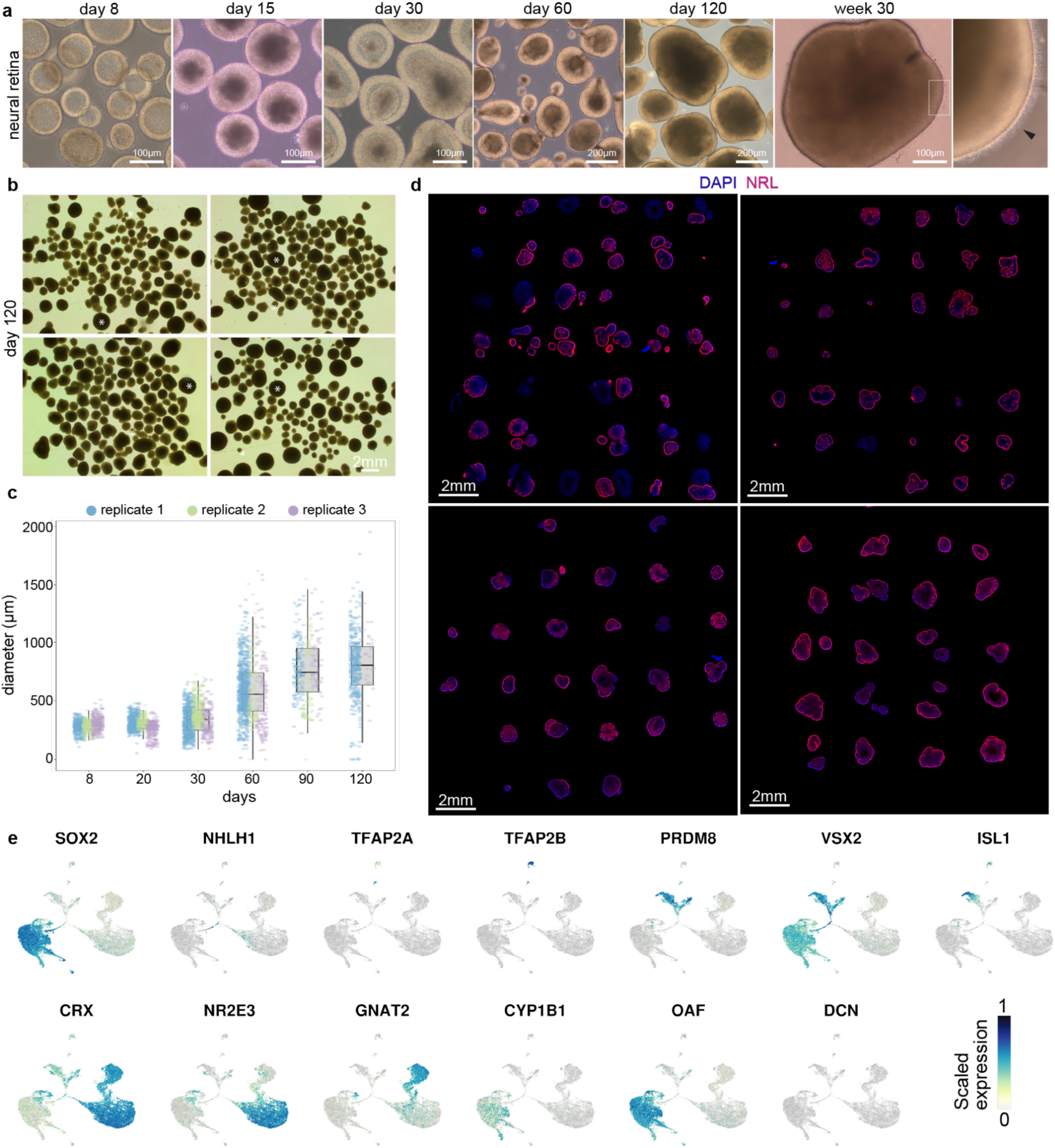
Retinal organoid generation from SLNs. **(a)** Brightfield images showing the morphological development of neural retina organoids over an extended culture period, from day 8 to week 30). Inset shows a magnified view of the retinal organoid periphery. Arrowhead marks photoreceptor outer segments. **(b)** Representative low magnification brightfield images illustrating selection of retinal organoids based on morphology. Dark and small organoids similar to ones marked by white stars were selected out. **(c)** Line graph showing the change in organoid diameter over time during differentiation. 2-4 replicates were quantified for each time point. **(d)** Representative confocal fluorescence microscopy images of retinal organoids from 4 independent replicates in a tissue microarray at day 120. Images show the spatial distribution of the rod photoreceptor marker NRL and nuclei (DAPI). **(e)** Feature plots showing expression of selected marker genes from cell types identified in figure 4l.

